# A robust benchmark for germline structural variant detection

**DOI:** 10.1101/664623

**Authors:** Justin M. Zook, Nancy F. Hansen, Nathan D. Olson, Lesley M. Chapman, James C. Mullikin, Chunlin Xiao, Stephen Sherry, Sergey Koren, Adam M. Phillippy, Paul C. Boutros, Sayed Mohammad E. Sahraeian, Vincent Huang, Alexandre Rouette, Noah Alexander, Christopher E. Mason, Iman Hajirasouliha, Camir Ricketts, Joyce Lee, Rick Tearle, Ian T. Fiddes, Alvaro Martinez Barrio, Jeremiah Wala, Andrew Carroll, Noushin Ghaffari, Oscar L. Rodriguez, Ali Bashir, Shaun Jackman, John J Farrell, Aaron M Wenger, Can Alkan, Arda Soylev, Michael C. Schatz, Shilpa Garg, George Church, Tobias Marschall, Ken Chen, Xian Fan, Adam C. English, Jeffrey A. Rosenfeld, Weichen Zhou, Ryan E. Mills, Jay M. Sage, Jennifer R. Davis, Michael D. Kaiser, John S. Oliver, Anthony P. Catalano, Mark JP Chaisson, Noah Spies, Fritz J. Sedlazeck, Marc Salit, the Genome in a Bottle Consortium

**Affiliations:** Material Measurement Laboratory, National Institute of Standards and Technology, 100 Bureau Dr, MS8312, Gaithersburg, MD 20899; National Human Genome Research Institute, National Institutes of Health, 5625 Fishers Lane, Rockville, MD 20852; National Center for Biotechnology Information, National Library of Medicine, National Institutes of Health, 45 Center Drive, Bethesda, MD, 20894; Department of Human Genetics, University of California, Los Angeles; Roche Sequencing Solutions, Belmont, CA, 94002, USA; Ontario Institute for Cancer Research, 661 University Ave, Suite 510, Toronto, ON M5G 0A3; Charles-Bruneau Cancer Centre, Division of Hematology-oncology, CHU Sainte-Justine, Montreal, Canada; Molecular Biology Institute, University of California, Los Angeles; Weill Cornell Medicine, 1300 York Ave., New York, NY 10065; Bionano Genomics, Inc. 9540 Towne Centre Drive, Ste. 100, San Diego, CA 92121; Davies Research Centre, School of Animal and Veterinary Sciences, University of Adelaide, Roseworthy SA 5371, Australia; 10x Genomics, Pleasanton, California 94566, USA; Broad Institute of Harvard and MIT, 415 Main Street, Cambridge, MA 02142; Google, 1600 Amphitheater Pkwy, Mountain View, CA 94040; Genomics and Bioinformatics, Texas A&M AgriLife Research, Texas A&M University, 1500 Research Parkway, Suite 250B, College Station, TX 77845; TAMU HPRC, Texas A&M University, Mail Stop 3361, College Station, TX 77843-3361; Department of Genetics and Data Sciences, Icahn School of Medicine at Mount Sinai, 1 Gustave L. Levy Place New York, NY 10029-5674; BC Cancer Genome Sciences Centre, 100-570 W 7th Ave, Vancouver, BC, V5Z 4S6, Canada; Biomedical Genetics, Dept of Medicine, Boston University Medical School, 72 East Concord Street, Boston MA 02118; Pacific Biosciences, Menlo Park, CA 94025, USA; Department of Computer Engineering, Bilkent University, Ankara 06800, Turkey; Department of Computer Engineering, Konya Food and Agriculture University, Konya 42080, Turkey; Departments of Computer Science and Biology, Johns Hopkins University, Baltimore, MD, 21218; Department of Genetics, Harvard Medical School, Boston, MA; Saarland University and Max Planck Institute for Informatics, Saarland Informatics Campus E2.1, 66123 Saarbrücken, Germany; Department of Bioinformatics and Computational Biology, MD Anderson Cancer Center, Houston, TX, 77030; Department of Computer Science, Rice University, Houston, TX, 77005; Bioinformatics R&D, Spiral Genetics, Seattle WA 98104; Rutgers Cancer Institute of New Jersey, New Brunswick, NJ, USA Department of Pathology, Robert Wood Johnson Medical School, New Brunswick, NJ, USA; Department of Computational Medicine and Bioinformatics, University of Michigan Medical School, 100 Washtenaw Avenue, Ann Arbor, MI 48109, USA; Nabsys 2.0, LLC, 60 Clifford St, Providence, RI 02903; Quantitative and Computational Biology, University of Southern California, 1050 Childs Way RRI 408H, Los Angeles, CA, 90089; Joint Initiative for Metrology in Biology, SLAC National Accelerator Lab, Stanford University, 435 Via Ortega, Stanford, CA 94305; Human Genome Sequencing Center, Baylor College of Medicine, One Baylor Plaza, Houston TX 77030

## Abstract

New technologies and analysis methods are enabling genomic structural variants (SVs) to be detected with ever-increasing accuracy, resolution, and comprehensiveness. Translating these methods to routine research and clinical practice requires robust benchmark sets. We developed the first benchmark set for identification of both false negative and false positive germline SVs, which complements recent efforts emphasizing increasingly comprehensive characterization of SVs. To create this benchmark for a broadly consented son in a Personal Genome Project trio with broadly available cells and DNA, the Genome in a Bottle (GIAB) Consortium integrated 19 sequence-resolved variant calling methods, both alignment- and *de novo* assembly-based, from short-, linked-, and long-read sequencing, as well as optical and electronic mapping. The final benchmark set contains 12745 isolated, sequence-resolved insertion and deletion calls ≥50 base pairs (bp) discovered by at least 2 technologies or 5 callsets, genotyped as heterozygous or homozygous variants by long reads. The Tier 1 benchmark regions, for which any extra calls are putative false positives, cover 2.66 Gbp and 9641 SVs supported by at least one diploid assembly. Support for SVs was assessed using *svviz* with short-, linked-, and long-read sequence data. In general, there was strong support from multiple technologies for the benchmark SVs, with 90 % of the Tier 1 SVs having support in reads from more than one technology. The Mendelian genotype error rate was 0.3 %, and genotype concordance with manual curation was >98.7 %. We demonstrate the utility of the benchmark set by showing it reliably identifies both false negatives and false positives in high-quality SV callsets from short-, linked-, and long-read sequencing and optical mapping.

## Introduction

Many diseases have been linked to structural variations (SVs), most often defined as genomic changes at least 50 base pairs (bp) in size, but SVs are challenging to detect accurately. Conditions linked to SVs include autism,^1^ schizophrenia, cardiovascular disease,^2^ Huntington’s Disease, and several other disorders.^3^ Far fewer SVs exist in germline genomes relative to small variants, but SVs affect more base pairs and each SV may be more likely to impact phenotype.^4–6^ While next generation sequencing technologies can detect many SVs, each technology and analysis method has different strengths and weaknesses. To enable the community to benchmark these methods, the Genome in a Bottle Consortium (GIAB) here developed benchmark SV calls and benchmark regions for the son (HG002/NA24385) in a broadly consented and available Ashkenazi Jewish trio from the Personal Genome Project,^7^ which are disseminated as National Institute of Standards and Technology (NIST) Reference Material 8392.^8,9^

Many approaches have been developed to detect SVs from different sequencing technologies. Microarrays can detect large deletions and duplications, but not with sequence-level resolution.^10^ Since short reads (<<1000bp) are often smaller than or similar to the SV size, bioinformaticians have developed a variety of methods to infer SVs, including using split reads, discordant read pairs, depth of coverage, and local *de novo* assembly. Linked reads add long-range (100kb+) information to short reads, enabling phasing of reads for haplotype-specific deletion detection, large SV detection,^11–13^ and diploid *de novo* assembly.^14^ Long reads (>>1000bp), which can fully traverse many more SVs, further enable SV detection, often sequence-resolved, using mapped reads,^15,16^ local assembly after phasing long reads,^6,17^ and global *de novo* assembly.^18,19^ Finally, optical mapping and electronic mapping provide an orthogonal approach capable of determining the approximate size and location of insertions, deletions, inversions, and translocations while spanning even very large SVs.^20–22^

GIAB recently published benchmark sets for small variants for seven genomes,^9,23^ and the Global Alliance for Genomics and Health Benchmarking Team established best practices for using these and other benchmark sets to benchmark germline variants.^24^ These benchmark sets are widely used in developing, optimizing, and demonstrating new technologies and bioinformatics methods, as well as part of clinical laboratory validation.^12,15,25,26^ Benchmarking tool development has also been critical to standardize definitions of performance metrics, robustly compare VCFs with different representations of complex variants, and enable stratification of performance by variant type and genome context. Benchmark set and benchmarking tool development is even more challenging and important for SVs given the wide spectrum of types and sizes of SVs, complexity of SVs (particularly in repetitive genome contexts), and that many SV callers output imprecise or imperfect breakpoints and sequence changes.

Several previous efforts have developed well-characterized SVs in human genomes. The 1000 Genomes Project catalogued copy-number variants (CNVs) and SVs in thousands of individuals.27,28 A subset of CNVs from NA12878 were confirmed and further refined to those with support from multiple technologies using SVClassify.^29^ The unique collection of Sanger sequencing from the HuRef sample has also been used to characterize SVs.^30,31^ Long reads were used to broadly characterize SVs in a haploid hydatidiform mole cell line.^32^ The Parliament framework was developed to integrate short and long reads for the HS1011 sample.^33^ Most recently, the Human Genome Structural Variation Consortium and the Genome Reference Consortium used short, linked, and long reads to develop phased, sequence-resolved SV callsets, greatly expanding the number of SVs in three trios from 1000 Genomes, particularly in tandem repeats.^6,34^ Detection of somatic SVs in cancer genomes is a very active field, with numerous methods in development.^35–37^ While some of the problems are similar between germline and somatic SV detection, somatic detection is complicated by the need to distinguish somatic from germline events in the face of differential coverage, subclonal mutations and impure tumor samples, amongst others.^38,39^

We build on these efforts by focusing on enabling anyone to assess both false negatives (FNs) AND false positives (FPs) for a well-defined set of sequence-resolved insertions and deletions ≥50 bp in specified genomic regions. We evaluate the utility of the benchmark for measuring precision and recall of diverse callsets from different technologies. While we include SVs only discovered by long reads, we exclude regions with more than one SV, as these regions are not handled by current SV comparison and benchmarking tools. We also cluster calls by their specific sequence, improving upon previous work that clustered loosely by position, overlap, or size; we address challenges in comparing calls with different representations in repetitive regions to enable the integration of a wide variety of sequence-resolved input callsets from different technologies.

## Results

### Candidate SV callsets differ by sequencing technology and analysis method

We generated 28 sequence-resolved candidate SV callsets from 19 variant calling methods from 4 sequencing technologies for the Ashkenazi son (HG002), as well as 20 callsets each from the parents HG003 and HG004. We integrated a total of 68 callsets, where we define a “callset” as the result of a particular variant calling method using data from one or more technologies for an individual. The variant calling methods included 3 small variant callers, 9 alignment-based SV callers, and 7 global *de novo* assembly-based SV callers. The technologies included short-read (Illumina and Complete Genomics), linked-read (10x Genomics), and long-read (PacBio) sequencing technologies as well as SV size estimates from optical (Bionano) and electronic (Nabsys) mapping.

Figure 1 shows the number of SVs overlapping between our sequence-resolved callsets from different variant calling methods and technologies for HG002. In general, the concordance for insertions is lower than the concordance for deletions, except among long-read callsets, mostly because current short read-based methods do not sequence-resolve large insertions. This highlights the importance of developing benchmark SV sets to identify which callset is correct when they disagree, and potentially when both are incorrect even when they agree.

**Figure 1:**
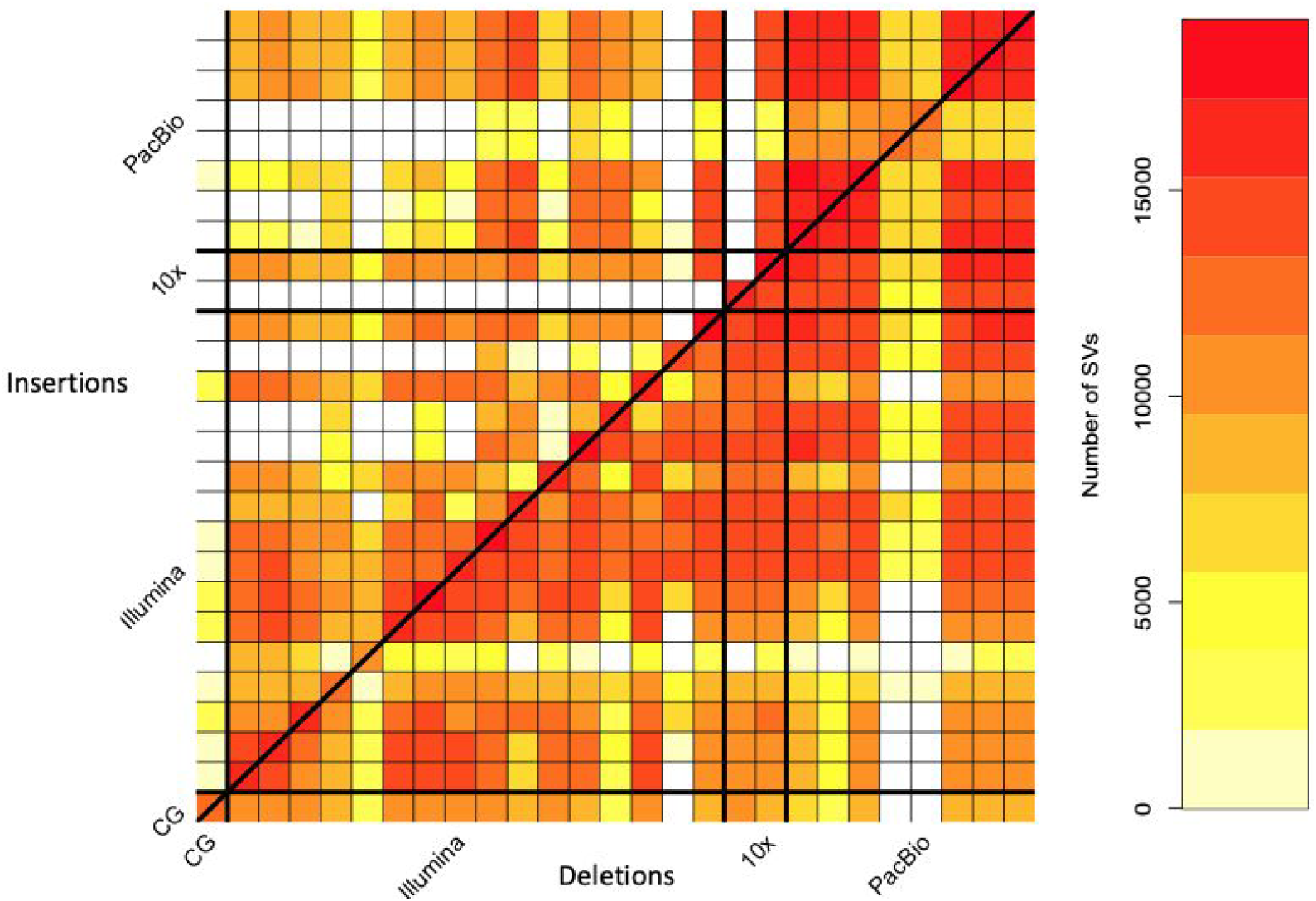
Pairwise comparison of sequence-resolved SV callsets obtained from multiple technologies and SV callers for SVs ≥50bp from HG002. Heatmap produced by SURVIVOR^40^ shows the number of SVs overlapping between the individual SV caller and technologies split between insertions (upper left) and deletions (lower right). The diagonal highlights the overall number of SVs per SV caller. Overall we obtained a quite diverse picture of SVs calls supported by each SV caller and technology, highlighting the need for benchmark sets.

### Design objectives for our benchmark SV set

Our objective was that, when comparing any callset (the “test set” or “query set”) to the “benchmark set,” it reliably identifies FPs and FNs. In practice, we aimed to demonstrate that most (ideally approaching 100%) of conflicts (both FPs *and* FNs) between any given test set and the benchmark set were actually errors in the test set. This goal is typically challenging to meet across the wide spectrum of sequencing technologies and calling methods. Secondarily, to the extent possible, our goal was for the benchmark set to include a large, representative variety of SVs in the human genome. By integrating results from a large suite of high-throughput, whole genome methods, each with their own signatures of bias, biases from any particular method are minimized. We systematically establish the “benchmark regions” in this genome in which we are close to comprehensively characterizing SVs. We exclude regions from our benchmark if we could not reliably reach near-comprehensive characterization (e.g., in segmental duplications). Importantly, we demonstrate the benchmark set is fit for purpose for benchmarking by presenting examples of comparisons of SVs from multiple technologies and manual curation of discordant calls.

### Benchmark set is formed by clustering and evaluating support for candidate SVs

We integrated all sequence-resolved candidate SV callsets to form the benchmark set, using the process described in **Figure 2**. Since candidate SV calls often differ in their exact breakpoints, size, and/or sequence change estimated, we used a new method called SVanalyzer (https://svanalyzer.readthedocs.io) to cluster calls estimating similar sequence changes. This new method was needed to account for both differences in SV representation (e.g., different alignments within a tandem repeat) and differences in the precise sequence change estimated. Of the 498876 candidate insertion and deletion calls ≥50 bp in the son-father-mother trio, 296761 were unique after removing duplicate calls and calls that were the same when taking into account representation differences (e.g., different alignment locations in a tandem repeat). When clustering variants for which the estimated sequence change was <20 % divergent, 128715 unique SVs remain. We then filtered to retain SV clusters supported by: more than one technology, ≥5 callsets from a single technology, Bionano, or Nabsys. The 30062 SVs remaining were then evaluated and genotyped in each member of the trio using svviz ^41^ to align reads to reference and alternate alleles from PCR-free Illumina, Illumina 6 kbp mate-pair, haplotype-partitioned 10x Genomics, and PacBio with and without haplotype partitioning. We further filtered for SVs covered in HG002 by 8 or more PacBio reads (mean coverage of about 60), with at least 25% of PacBio reads supporting the alternate allele and consistent genotypes from all technologies that could be confidently assessed with svviz. This left 19748 SVs. The number of PacBio reads supporting the SV allele and reference allele for each benchmark SV is in Supplementary Figure 1.

**Figure 2:**
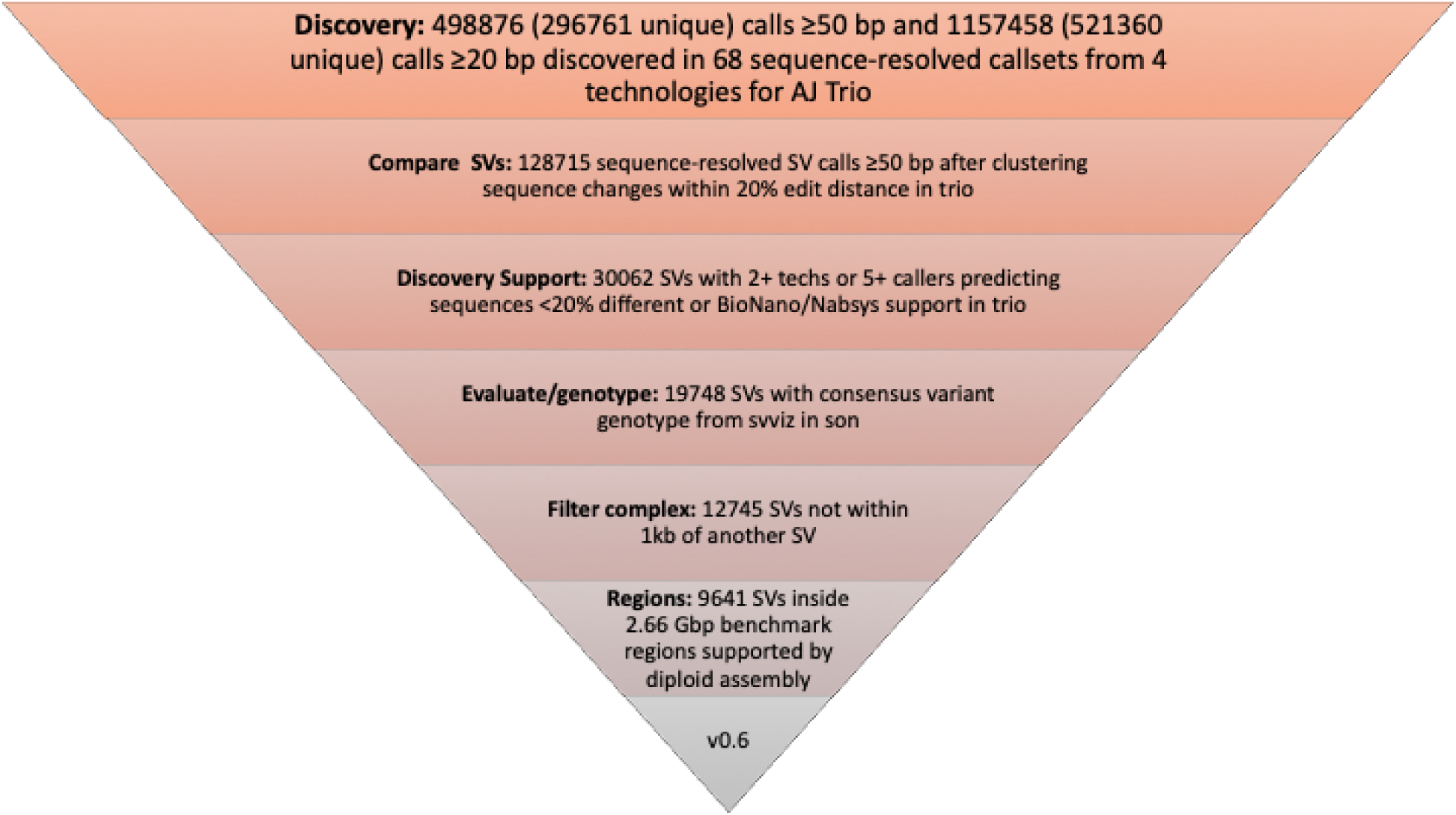
Process to integrate SV callsets from different technologies and analysis methods and form the benchmark set. Approximately 0.5 million input SV calls were locally clustered based on their estimated sequence change, and we kept only those discovered by at least two technologies or at least 5 callsets in the trio. We then used svviz with short, linked, and long reads to evaluate and genotype these calls, keeping only those with a consensus heterozygous or homozygous variant genotype in the son. We filtered potentially complex calls in regions with multiple discordant SV calls, as well as regions around 20 bp to 49 bp indels, and our final Tier 1 benchmark set included 12745 total insertions and deletions ≥50 with 9641 inside the 2.66 Gbp of the genome where diploid assemblies had no additional SVs beyond those in our benchmark set. We also define a Tier 2 set of 6007 additional regions where there was substantial support for one or more SVs but the precise SV was not yet determined.

In our evaluations of these well-supported SVs, we found that 12745 were isolated, while 7003 (35 %) were within 1000 bp of another well-supported SV call. Upon manual curation, we found that the variants within 1000 bp of another variant were mostly in tandem repeats and fell into several classes: (1) true complex variants with more than one SV call on the same haplotype, (2) true compound heterozygous variant with different SV calls on each haplotype, and (3) regions where some methods had the correct SV call and others had inaccurate sequence, size, or breakpoint estimates, but svviz still aligned reads to it because reads matched it better than the reference. We chose to exclude these clustered SVs from our benchmark set because methods do not exist to confidently distinguish between the above classes, nor do SV comparison tools for robust benchmarking of complex and compound structural variants.

Finally, to enable assessment of both FNs and FPs, benchmark regions were defined using diploid assemblies and candidate variants. These regions were designed such that our benchmark variant callset should contain almost all true SVs within these regions. These regions define our Tier 1 benchmark set, which spans 2.66 Gbp and includes 9641 benchmark SVs. These regions exclude 1837 of the 12745 SVs because they were within 50 bp of a 20 bp to 49 bp indel; they exclude an additional 856 SVs within 50 bp of a candidate SV for which no consensus genotype could be determined; and they exclude an additional 411 calls that were not fully supported by a diploid assembly as the only SV in the region. A large number of annotations are associated with the Tier 1 SV calls (e.g., number of discovery callsets from each technology, number of reads supporting reference and alternate alleles from each technology, number of callsets with exactly matching sequence estimates), which enable users to filter to a more specific callset. We also define Tier 2 regions that delineate 6007 additional regions in addition to the 12745 isolated SVs, which are regions with substantial evidence for one or more SVs but we could not precisely determine the SV. For the Tier 2 regions, multiple SVs within 1 kb or in the same or adjacent tandem repeats are counted as a single region, so many SV callers would be expected to call more than 6007 SVs in these regions.

### Benchmark calls are well-supported

The 12745 isolated SV calls had size distributions consistent with previous work detecting SVs from long reads,^6,15,17,26^ with the clear, expected peaks for Alu insertions and deletions near 300 bp and for full-length LINE1 insertions and deletions near 6000 bp (**Figure 3**). SVs have an exponentially decreasing abundance vs. size if they fall in tandem repeats longer than 100 bp in the reference. Interestingly, there are more large insertions than large deletions in tandem repeats, despite insertions being more challenging to detect. This is consistent with previous work detecting SVs from long read sequencing^15,17^ and may result from instability of tandem repeats in the BAC clones used to create the reference genome.^42^

**Figure 3:**
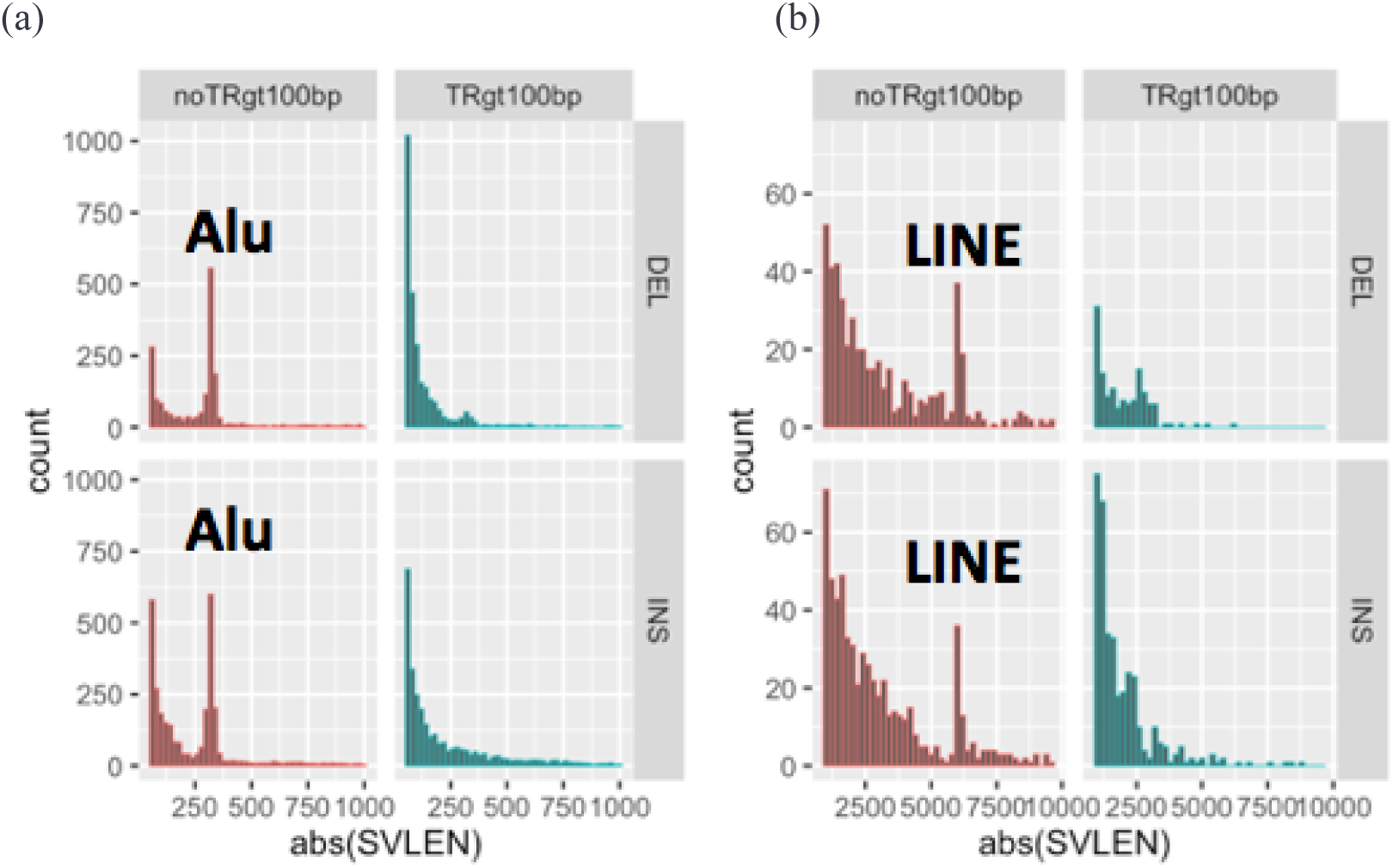
Size distributions of deletions and insertions in the benchmark set. Distributions shown on separate scales for (a) 50 bp to 1000 bp and (b) 1000 bp to 10000 bp. Variants are stratified into deletions (DEL) and insertions (INS), and into SVs overlapping (TRgt100bp) and not overlapping (noTRgt100bp) tandem repeats longer than 100bp in the reference. The expected peaks near 300 bp are labeled as Alu mobile elements and near 6000 bp are labeled as LINE mobile elements.

When evaluating the support for our benchmark SVs, approximately 50 % of long reads more closely matched the SV allele for heterozygous SVs, and approximately 100 % for homozygous SVs, as expected (**Figure 4**). While short reads clearly supported and differentiated homozygous and heterozygous genotypes for many SVs, the support for heterozygous calls was less balanced, with a mode around 30%, and they did not definitively genotype 35 % of deletions and 47 % of insertions in tandem repeats because reads were not sufficiently long to traverse the repeat. These results highlight the difficulty in detecting SVs with short reads in long tandem repeats, as a sizeable fraction of reads containing the variant either map without showing the variant or fail to map at all. We also found high concordance with Bionano, with 90 % of our sequence-resolved SVs > 1000 bp having a size within 22 % of the size estimated by Bionano, and 75 % having a size with 7 % of Bionano. In general, there was strong support from multiple technologies for the benchmark SVs, with 90 % of the Tier 1 SVs having support from more than one technology.

**Figure 4:**
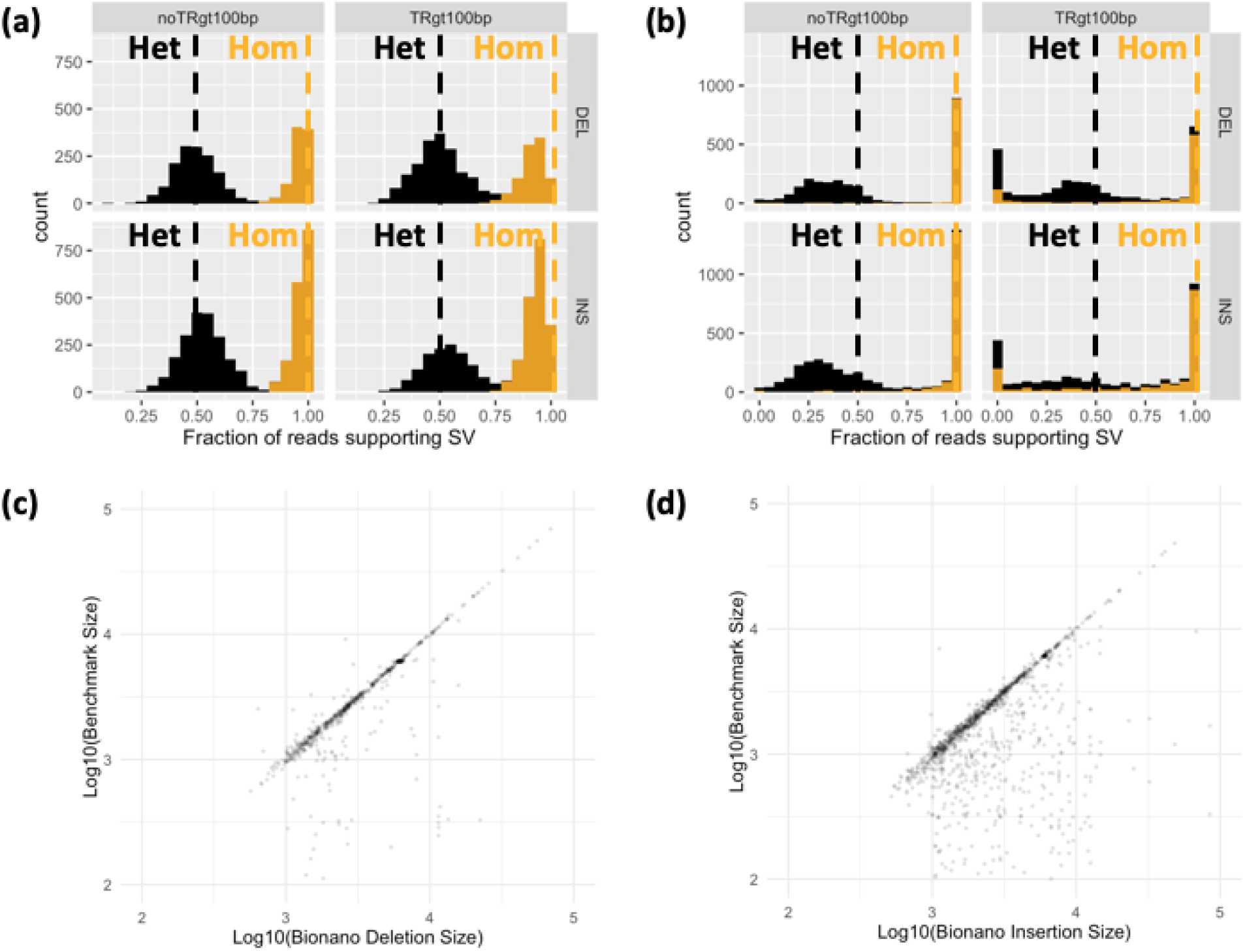
Support for benchmark SVs by long reads, short reads, and optical mapping. Histograms show the fraction of (a) PacBio and (b) Illumina 150 bp reads that aligned better to the SV allele than to the reference allele using svviz, colored by v0.6 genotype, where black is heterozygous and orange is homozygous. Variants are stratified into deletions (DEL) and insertions (INS), and into SVs overlapping (TRgt100bp) and not overlapping (noTRgt100bp) tandem repeats longer than 100bp in the reference. Vertical dashed lines correspond to the expected fractions 0.5 for heterozygous and 1.0 for homozygous variants. In (c) and (d), the sequence-resolved size in the v0.6 benchmark set is plotted against the size estimated by BioNano in any overlapping intervals, where points below the diagonal represent smaller sequence-resolved SVs in the overlapping interval.

For SVs on autosomes, we also identified if genotypes were consistent with Mendelian inheritance. When limiting to 7973 autosomal SVs in the benchmark set for which a consensus genotype from svviz was determined for both of the parents, only 20 violated Mendelian inheritance. Upon manual curation of these 20 sites, 16 were correct in HG002 (mostly misidentified as homozygous reference in both parents due to lower long read sequencing coverage), 1 was a likely de novo deletion in HG002 (17:51417826-51417932), 1 was a deletion in the T cell receptor alpha locus known to undergo somatic rearrangement (14:22918114-22982920), and 2 were insertions mis-genotyped as heterozygous in HG002 when in fact they were likely homozygous variant or complex (2:232734665 and 8:43034905). Supplementary Table 1 is a detailed contingency table of genotypes in the son, father, and mother.

**Table 1:** Heuristics for genotype determination from each dataset. Svviz was used to determine the genotype for HG002 using different heuristics for each dataset. The cut-offs for weighted alternate (ALT) and reference (REF) counts were determined manually from looking at distributions for different size ranges. For each technology, genotypes were determined based on the proportion of reads supporting the SV. If the ALT and REF counts did not meet the criteria in this table for a particular dataset, the genotype was considered uncertain. In addition, the genotype was considered uncertain for PacBio if it overlapped a tandem repeat longer than 10 kbp, for Illumina and 10x if it overlapped a tandem repeat longer than 100 bp, and for all datasets if it overlapped a segmental duplication >10 kbp.

| Dataset(s) | Minimum Coverage | Proportion of reads supporting SV ($x = \text{ALT}/(\text{REF}+\text{ALT})$) | Genotype Label |
| --- | --- | --- | --- |
| PacBio | $\text{ALT}+\text{REF} \geq 8$ | $x < 0.1$ | 0/0 |
| | | $0.25 < x < 0.75$ | 0/1 |
| | | $x > 0.9$ | 1/1 |
| Illumina 250bp and Illumina Mate-Pair | $\text{ALT}+\text{REF} \geq 8$ | $x < 0.05$ | 0/0 |
| | | $0.1 < x < 0.9$ | 0/1 |
| | | $x > 0.95$ | 1/1 |
| Haplotype-partitioned 10x Genomics and PacBio (haplotype in subscript) | $\text{ALT}_1 + \text{REF}_1 \geq 5$<br>AND<br>$\text{ALT}_2 + \text{REF}_2 \geq 5$ | $x_1 < 0.05$ AND $x_2 < 0.05$ | 0/0 |
| | | $(x_1 > 0.95$ AND $x_2 < 0.05)$ OR<br>$(x_2 > 0.95$ AND $x_1 < 0.05)$ | 0/1 |
| | | $x_1 > 0.95$ AND $x_2 > 0.95$ | 1/1 |

The GIAB community also manually curated a random subset of SVs from different size ranges in the union of all discovered SVs.^43^ When comparing the consensus genotype from expert manual curation to our benchmark SV genotypes, 627/635 genotypes agreed. Most discordant genotypes were identified as complex by the curators, with a 20 bp to 49 bp indel near an SV in our benchmark set, because they were asked to include indels 20 bp to 49 bp in size in their curation, whereas our SV benchmark set focused on SVs >49 bp.

### Benchmark set is useful for identifying false positives and false negatives across technologies

Our goal in designing this SV benchmark set was that, when comparing any callset to our benchmark VCF within the benchmark BED file, most putative FPs and FNs should be errors in the tested callset. To determine if we meet this goal, we benchmarked several callsets from assembly- and non-assembly-based methods that use short or long reads. We developed a new benchmarking tool *truvari* (*https://github.com/spiralgenetics/truvari*) to perform these comparisons at a matching stringency requiring the variant size to be within 30 % of the benchmark size, and the position to be within 2kb. Truvari enables users to specify matching stringency for size, sequence, and/or distance. An alternative benchmarking tool developed more recently, which has more sophisticated sequence matching, is *SVanalyzer SVbenchmark* (https://github.com/nhansen/SVanalyzer/blob/master/docs/svbenchmark.rst).

Upon manual curation of a random 10 FP and FN insertions and deletions (40 total SVs) from each callset being compared to the benchmark, nearly all of the FPs and FNs were errors in each of the tested callsets and not errors in the GIAB callset (**Figure 5**). The version of the *truvari* tool we used could not always account for all differences in representation, so if manual curation determined both the benchmark and test sets were correct, they were counted as correct. The only notable exception to the high GIAB callset accuracy was for FP insertions from the PacBio caller pbsv (https://github.com/PacificBiosciences/pbsv), for which about half of the putative FP insertions were true insertions missed in the benchmark regions.

**Figure 5:**
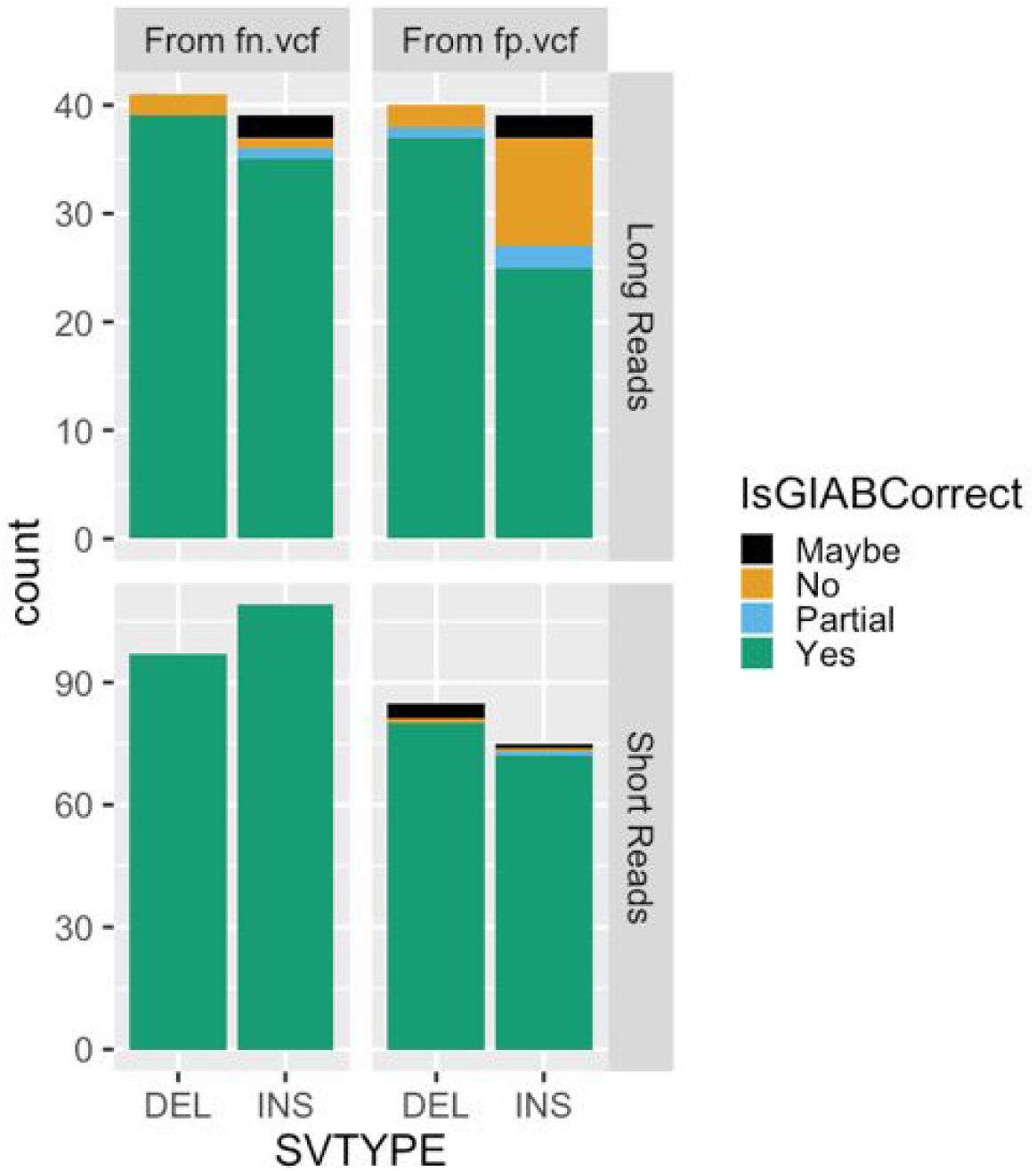
Summary of manual curation of putative FPs and FNs when benchmarking short and long reads against the v0.6 benchmark set. Most FP and FN SVs were determined to be correct in the v0.6 benchmark (green), but some were partially correct due to missing part of the SV in the region (blue), were incorrect in v0.6 (orange), or were in difficult locations where the evidence was unclear (black).

This suggests the GIAB callset may be missing approximately 5 % of true insertions in the benchmark regions. When comparing BioNano calls to our benchmark, we also found one region with multiple insertions where our benchmark had a heterozygous 1412 bp insertion at chr6:65000859, but we incorrectly called a homozygous 101 bp insertion in a nearby tandem repeat at chr6:65005337, when in fact there is an insertion of approximately 5400 bp in this tandem repeat on the same haplotype as the 1412 bp insertion, and the 101 bp insertion is on the other haplotype.

### Technologies and variant callers have different strengths and weaknesses

Amongst the extensive candidate SV callsets we collected from different technologies and analyses, we found that certain SV types and sizes in our benchmark set were discovered by fewer methods (**Figure 6**). In particular, more methods discovered sequence-resolved deletions than insertions, more methods discovered SVs not in tandem repeats, and the most methods discovered deletions smaller than 1000 bp not in tandem repeats. These results confirm the intuition that SV detection outside of repeats is simpler than within repeats, and that deletions are simpler to detect than insertions since deletions do not require mapping to new sequence. **Figure 7** further shows that the fewest SVs were missed by the union of all long read discovery methods. The only exception was (50 to 99) bp deletions, which were all found by at least one short read discovery method. Many insertions >300 bp that were not discovered by any short read method could be accurately genotyped in this sample by short reads. Interestingly, many deletions and insertions <300 bp that were not genotyped accurately by short reads were discovered by at least one short read-based method. This likely reflects a limitation of the heuristics we used for genotyping, which reduces the false positive rate but may increase the false negative rate. Both discovery and genotyping based on short reads had limitations for SVs in tandem repeats. These results confirm the importance of long read data for comprehensive SV detection.

**Figure 6:**
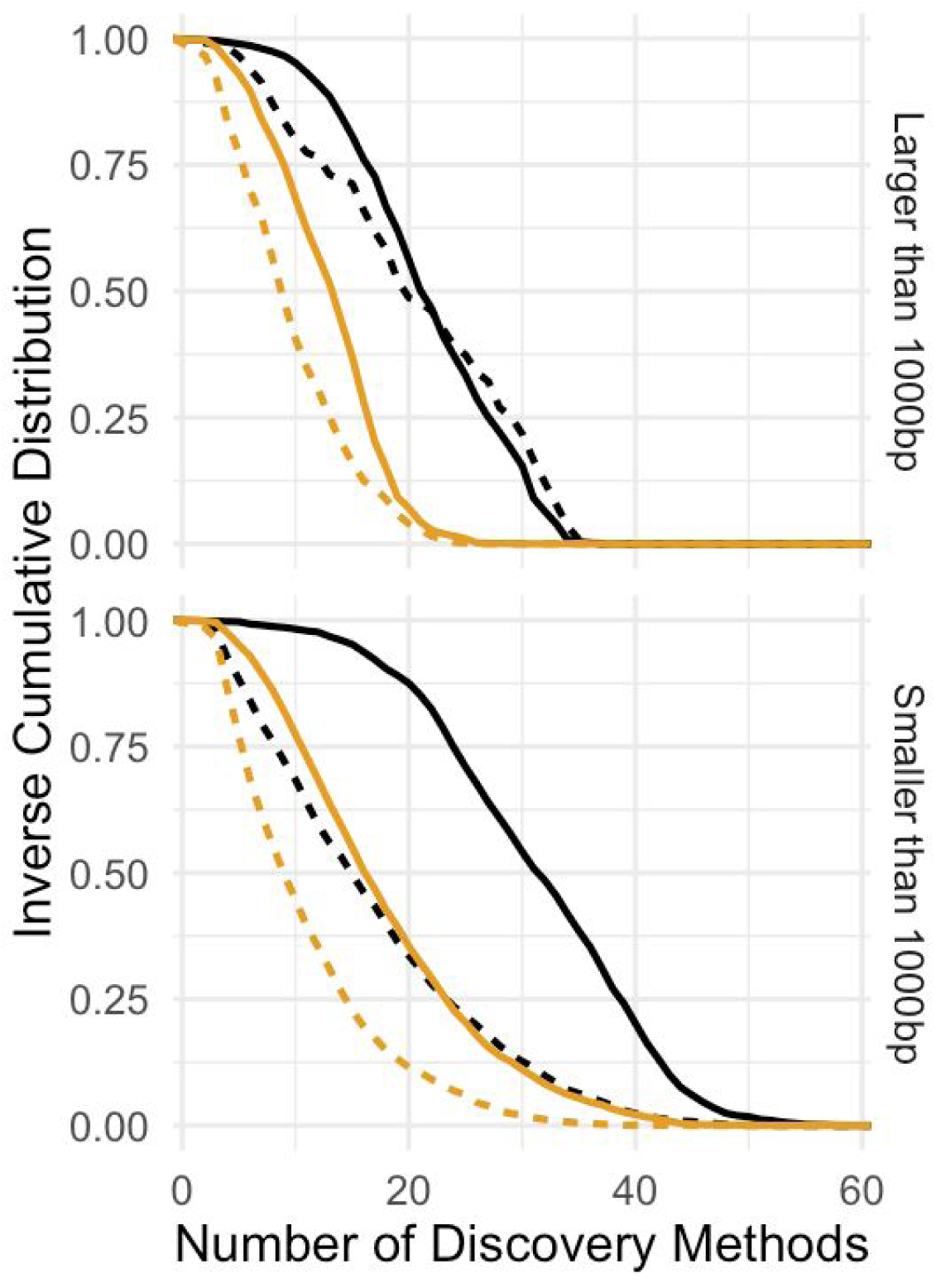
Inverse cumulative distribution showing the number of discovery methods that supported each SV. All 68 callsets from all variant calling methods and technologies in all three members of the trio are included in these distributions. SVs larger than 1000 bp (top) are displayed separately from SVs smaller than 1000 bp (bottom). Results are stratified into deletions (black) and insertions (orange), and into SVs overlapping (solid) and not overlapping (dashed) tandem repeats longer than 100 bp in the reference.

**Figure 7:**
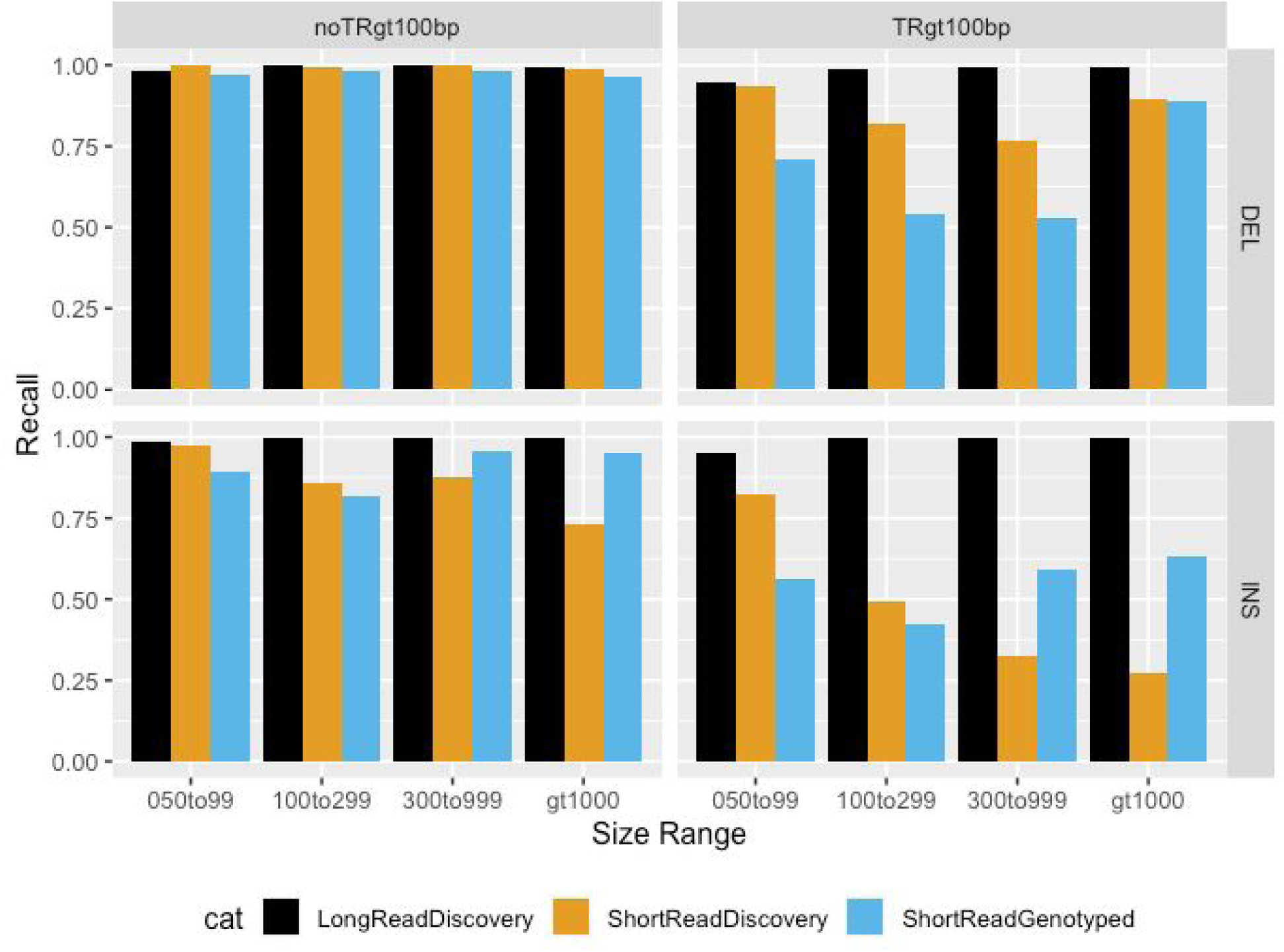
Comparison of false negative rates for the union of all long read-based SV discovery methods, the union of all short read-based discovery methods, and paired-end and mate-pair short read genotyping of known SVs. Variants are stratified into deletions (DEL) and insertions (INS), and into SVs overlapping (TRgt100bp) and not overlapping (noTRgt100bp) tandem repeats longer than 100 bp in the reference. SVs are also stratified by size into 50 bp to 99 bp (050to99), 100 bp to 299 bp (100to299), 300 bp to 999 bp (300to999), and ≥1000 bp (gt1000).

### Sequence-resolved benchmark calls have annotations related to base-level accuracy

We provide sequence-resolved calls in our benchmark set to enable benchmarking of sequence change predictions, but importantly not all calls are perfect on a base-level. When discovered SVs from multiple callsets have exactly matching sequence changes, we output the sequence change from the largest number of callsets. However, as shown in **Figure 8**, not all benchmark SVs have calls that exactly matched between discovery callsets. For deletions not in tandem repeats, at least 99 % of the calls had exact matches, but there were no exact matches for ~5% of DELs in TRs, and for large insertions no exact matches existed for ~50% of the calls. This is likely because SVs in tandem repeats and larger insertions are more likely to be discovered only by methods using relatively noisy long reads.

**Figure 8:**
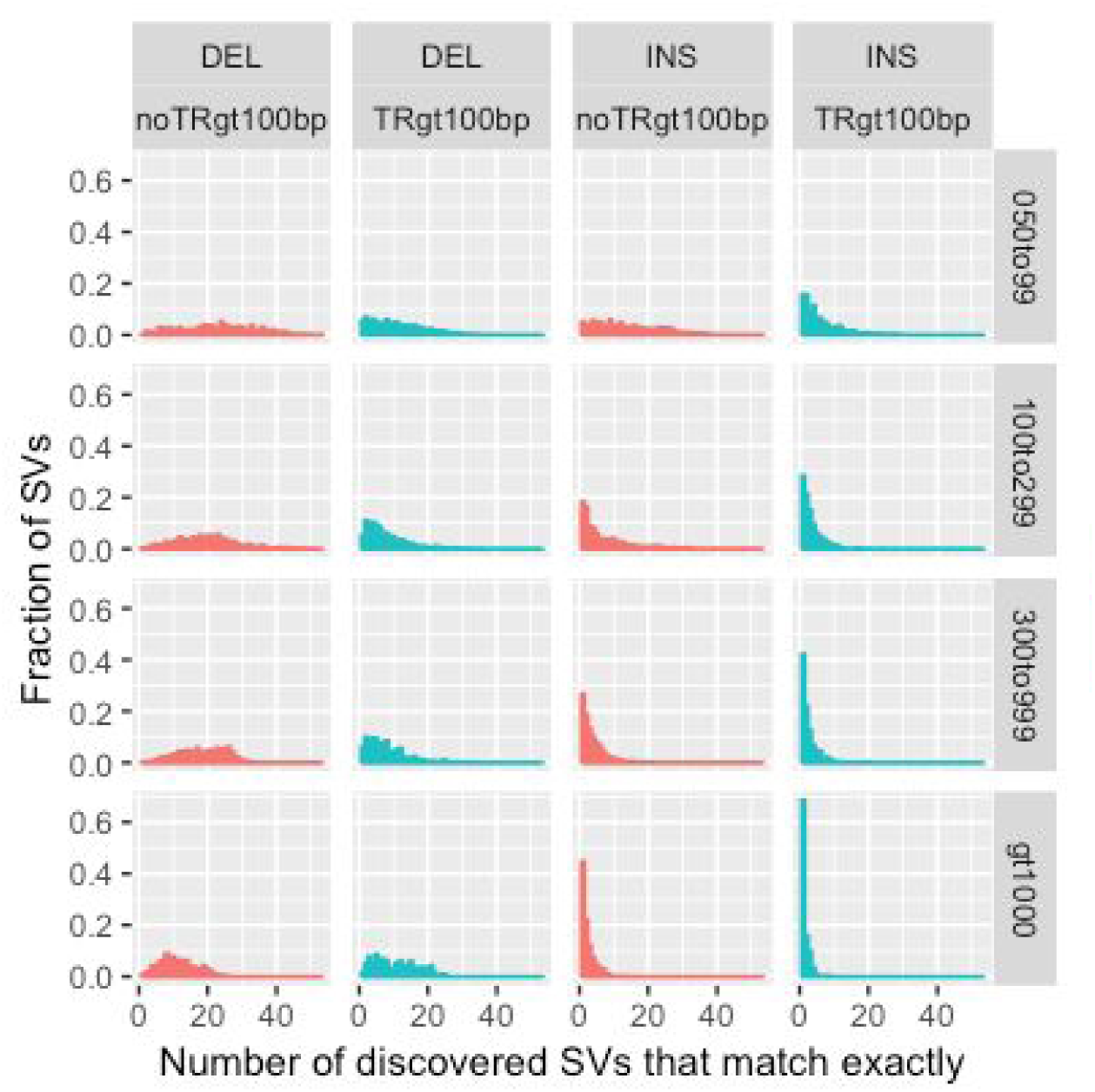
Fraction of SVs for each number of discovery callsets that estimated exactly matching sequence changes. Variants are stratified into deletions (DEL) and insertions (INS), and into SVs overlapping (TRgt100bp) and not overlapping (noTRgt100bp) tandem repeats longer than 100 bp in the reference. SVs are also stratified by size into 50 bp to 99 bp (050to99), 100 bp to 299 bp (100to299), 300 bp to 999 bp (300to999), and ≥1000 bp (gt1000).

## Discussion

We have integrated sequence-resolved SV calls from diverse technologies and SV calling approaches to produce a new benchmark set enabling anyone to assess both FN and FP rates. This benchmark is useful for evaluating accuracy of SVs from essentially any genomic technology, including short, linked, and long read sequencing technologies, optical mapping and electronic mapping. This resource of benchmark SVs, data from a variety of technologies, and SVs from a variety of methods are all publicly available without embargo, and we encourage the community to give feedback and participate in GIAB to continue to improve and expand this benchmark set in the future.

When developing this benchmark set, several trade-offs were made. Most notably, we chose to exclude complex SVs and SVs for which we could not determine a consensus sequence. Limiting our set to isolated insertions and deletions removed approximately one half of SVs for which there was strong support that some SV occurred. However, by excluding these complex regions from our SV benchmark set, it enables anyone to use our sequence comparison-based benchmarking tools to confidently and automatically identify FPs and FNs at different matching stringencies (*e.g*., matching based on SV sequence, size, type, and/or genotype). Bionano also identified large heterozygous events outside the benchmark regions, and future work will be needed to sequence-resolve these complex events, often near segmental duplications. In addition to our standard Tier 1 benchmark set, we also provide a set of Tier 2 regions in which we found substantial evidence for an SV but it was complex or we could not determine the precise SV. We also exclude regions from our benchmark set around putative indels (20 to 49) bp in size, which minimizes unreliable putative FP and FN SVs around clustered indels or variants just under or above 50 bp.

Our benchmark also currently does not include more complicated forms of structural variations including inversions, duplications (except for calls annotated as tandem duplications), very large copy number variants (only one deletion and one insertion >100 kb), calls in segmental duplications, calls in tandem repeats >10 kbp, or translocations. We also do not explicitly call duplications, though in practice our insertions frequently are tandem duplications, and we have provisionally labeled them as such using SVanalyzer svwiden in the REPTYPE annotation in the benchmark VCF. Future work in GIAB will use new technologies and analysis methods to include new SV types and more challenging SVs. When using our current benchmark, it is critical to understand it does not enable performance assessment for all SV types nor the most challenging SVs.

GIAB is currently collecting new candidate SV callsets for GRCh37 and GRCh38 from new data types (e.g., PacBio Circular Consensus Sequencing^26^ and Oxford Nanopore ultra long reads^44^), new and updated SV callers, and new diploid *de novo* assemblies. We are also refining the integration methods (e.g., to include inversions), and developing an integration pipeline that is easier to reproduce. In the next several months, we plan to release improved benchmark sets for GRCh37 and GRCh38 using these new methods similar to how we have maintained and updated the small variant callsets for these samples over time. We will also use the reproducible integration pipeline developed here to benchmark SVs for all 7 GIAB genomes. We will continue to refine these methods to access more difficult SVs in more difficult regions of the genome. Finally, we plan to develop a manuscript describing best practices for using this benchmark set to benchmark any other SV callset, similar to our recent publication for small variants,^24^ with refined SV comparison tools and standardized definitions of performance metrics.

## Data availability

Raw sequence data were previously published in Scientific Data (DOI: 10.1038/sdata.2016.25), and were deposited in the NCBI SRA with the accession codes SRX847862 to SRX848317, SRX1388732 to SRX1388743, SRX852933, SRX5527202, SRX5327410, and SRX1033793-SRX1033798. 10x Genomics Chromium bam files used are at ftp://ftp-trace.ncbi.nlm.nih.gov/giab/ftp/data/AshkenazimTrio/analysis/10XGenomics_ChromiumGenome_LongRanger2.2_Supernova2.0.1_04122018/. The data used in this manuscript and other datasets for these genomes are available in ftp://ftp-trace.ncbi.nlm.nih.gov/giab/ftp/data/, and in the NCBI BioProject PRJNA200694.

The v0.6 SV benchmark set (***only compare to variants in the Tier 1 vcf inside the Tier 1 bed with the FILTER “PASS”***) for HG002 on GRCh37 is available at: ftp://ftp-trace.ncbi.nlm.nih.gov/giab/ftp/data/AshkenazimTrio/analysis/NIST_SVs_Integration_v0.6/

Input SV callsets, assemblies, and other analyses for this trio are available under: ftp://ftp-trace.ncbi.nlm.nih.gov/giab/ftp/data/AshkenazimTrio/analysis/

## Acknowledgments

We thank many Genome in a Bottle Consortium Analysis Team members for helpful discussions about the design of this benchmark. Certain commercial equipment, instruments, or materials are identified to specify adequately experimental conditions or reported results. Such identification does not imply recommendation or endorsement by the National Institute of Standards and Technology, nor does it imply that the equipment, instruments, or materials identified are necessarily the best available for the purpose. Chunlin Xiao and Steve Sherry were supported by the Intramural Research Program of the National Library of Medicine, National Institutes of Health.

## Methods

### Code availability

Scripts for integrating candidate structural variants to form the benchmark set in this manuscript are available in a GitHub repository at https://github.com/jzook/genome-data-integration/tree/master/StructuralVariants/NISTv0.6. Publicly available software used to generate input callsets is described below in the methods.

### Tier 1 Benchmark Integration process

The GIAB v0.6 Tier 1 Benchmark SV Set was generated using the following heuristics from the union vcf generated from the discovery callsets described below (68 callsets from 19 variant callers and 4 technologies for the GIAB Ashkenazi trio) at ftp://ftp-trace.ncbi.nlm.nih.gov/giab/ftp/data/AshkenazimTrio/analysis/NIST_UnionSVs_12122017/union_171212_refalt.sort.vcf.gz

1. Sequence-resolved variants with at least 20% sequence similarity were merged into a single vcf line using the SVanalyzer merge command (https://github.com/nhansen/SVanalyzer). SVanalyzer merges variants by aligning and comparing their extended alternate haplotypes rather than by using size and overlap rules. Pairs of variants whose alternate haplotypes have normalized edit distance and size difference less than or equal to 20% of the length of the extended haplotype region are considered to be matches and clustered into single variants. See the section “Clustering of sequence-resolved variants with SVmerge” for a more detailed description.
2. Variants supported by at least two technologies (including BioNano and Nabsys) or by at least 5 callsets from a single technology had evidence for them evaluated and were genotyped using svviz2 with the four datasets in Table 1. Genotypes from Illumina and 10x were excluded in tandem repeats >100 bp in length, and genotypes from PacBio were excluded in tandem repeats >10000 bp. Genotypes from all datasets were excluded in segmental duplications >10000 bp. If the genotypes from all remaining datasets were concordant, and PacBio supported a genotype of heterozygous or homozygous variant, then the variant was included in downstream analyses.
3. If two or more supported variants ≥50 bp were within 1000 bp of each other, they were excluded because they are potentially complex or inaccurate.

In addition, benchmark regions were formed with the following process:

1. Use SVRefine to call variants from 3 PacBio-based (MsPAC, Phased-SV, and Falcon-unzip) and 1 10x-based (supernova) assemblies (results and methods at ftp://ftp-trace.ncbi.nlm.nih.gov/giab/ftp/data/AshkenazimTrio/analysis/NHGRI_SVrefine_04122018/)
2. Use SVRefine to compare variants from each assembly to our v0.6 PASS calls for HG2 allowing them to be up to 20 % different in all 3 distance measures, and only keep variants not matching a v0.6 call.
  a. Cluster the remaining variants from all assemblies and keep any that are supported by at least one long read assembly
3. Find regions for each assembly that are covered by exactly one contig for each haplotype (uses bedgraphs at link in #1)
4. Find the number of assemblies for which both haplotypes cover each region
5. Subtract regions around variants remaining after #2, using svwiden’s repeat-expanded coordinates, and expanded further to include any overlapping repetitive regions from Tandem Repeat Finder, RepeatMasker SimpleRepeats, and RepeatMasker LowComplexity, plus 50 bp on each end.
6. High confidence regions are regions in #4 covered by at least 1 assembly minus the regions in #5.
7. Further exclude any regions in the Tier 2 bed file of unresolved and clusters of variants, unless the Tier 2 region overlaps a Tier 1 PASS call.

### Tier 2 Benchmark Integration Process

We designed the draft Tier 2 benchmark set as a less conservative set of regions in which there appeared to be good evidence for at least one SV, but there were multiple SVs within 1 kb, multiple SVs within a tandem repeat, and/or different SV callers had different results for reasons that were not yet resolved. The process for forming the Tier 2 regions was:

1. Add 1000 bp to each side of any variants with the FILTER “ClusteredCalls” or “MaxEditDistgt04” and merge regions separated by <50 bp. Expand these regions to completely encompass any overlapping tandem repeats (after merging tandem repeats within 50 bp and adding 5 bp on each side, available at https://github.com/jzook/genome-data-integration/blob/master/StructuralVariants/NISTv0.6/repeats_trfSimplerepLowcomplex_merged50_slop5.bed.gz)
2. Take any variants ≥50 bp with the FILTER “NoConsensusGT”, expand these regions to completely encompass any overlapping tandem repeats (after merging tandem repeats within 50bp and adding 5 bp on each side), merge regions separated by <50 bp, and add 50 bp to each side.
3. After removing variants discovered by at least 2 technologies or 5 callsets (the inverse of those tested in the Tier 1 process above), cluster variants within 1000 bp (without considering type or sequence change), and find regions with clusters having calls from at least 2 technologies or 5 callsets. Expand these regions to completely encompass any overlapping tandem repeats (after merging tandem repeats within 50bp and adding 5 bp on each side).
4. Remove any regions from #1 and #2 that have any overlap with a Tier 1 benchmark call, and take the union of the resulting regions and the regions from #3. Merge regions within 50bp, and the result is the Tier 2 bed.

### Clustering of sequence-resolved variants with SVmerge

Structural variants are frequently flanked by stretches of repeated sequence which obscure the true position of the structural event. For this reason, we used a repeat-aware method to compare sequence-resolved structural variants, rather than relying on size and overlap-based rules. The program SVmerge, part of the SVanalyzer package (http://github.com/nhansen/SVanalyzer) was used to compare pairs of non-identical structural variant calls and cluster them based on distance measures. To calculate these measures of distance, SVmerge constructs alternate haplotype sequences corresponding to a common, widened region of the reference which includes all bases altered by either of the two variants. The two resulting alternate haplotypes are then compared by global alignment (Needleman Wunsch, as implemented in the the edlib software library),^45^ and the resulting alignment is used to calculate three normalized measures of difference: (1) the edit distance between the two alternate haplotypes, (2) the size difference in base pairs between the two alternate haplotypes, and (3) the maximum shift of coordinates in the global alignment between the two haplotypes. Each of these distances is then normalized by dividing by the mean length of the longer allele (reference or alternate) for each of the two variants.

To combine the structural variant calls into clusters, SVmerge creates an undirected graph in which variant calls are nodes and edges exist between pairs of calls having all three distances less than or equal to specified maximum values. Variants are then merged into a single cluster if they are within the same connected component of the resulting graph.

### Trio+linked read phased vcf and haplotype-partitioned bam files

To produce a chromosome-length phasing of small variants for the Ashkenazim trio, we combined variant calls from Real Time Genomics (ftp://ftp-trace.ncbi.nlm.nih.gov/giab/ftp/data/AshkenazimTrio/analysis/Rutgers_IlluminaHiSeq300X_rtg_11052015/rtg_allCallsV2.vcf.gz) with phased blocks produced by 10x Genomics in the following vcfs: ftp://ftp-trace.ncbi.nlm.nih.gov/giab/ftp/data/AshkenazimTrio/analysis/10XGenomics_ChromiumGenome_LongRanger2.1_09302016/NA24143_hg19/NA24143_hg19_phased_variants.vcf.gz ftp://ftp-trace.ncbi.nlm.nih.gov/giab/ftp/data/AshkenazimTrio/analysis/10XGenomics_ChromiumGenome_LongRanger2.1_09302016/NA24149_hg19/NA24149_hg19_phased_variants.vcf.gz ftp://ftp-trace.ncbi.nlm.nih.gov/giab/ftp/data/AshkenazimTrio/analysis/10XGenomics_ChromiumGenome_LongRanger2.1_09302016/NA24385_hg19/NA24385_hg19_phased_variants.vcf.gz

The single sample 10x Genomics VCF files were combined into multi-sample VCF using bcftools and all VCFs were split by chromosome (to facilitate easy parallelization with Snakemake). Then, WhatsHap (version 0.15+14.ga105b78)^46^ was used in pedigree-aware mode^47^ using the following command line:

whatshap phase --ped AJ.ped --indels --reference hg19.fasta rtg.vcf 10x-merged.vcf | bgzip > output.vcf

This vcf with whatshap haplotag was used to partition reads in the PacBio bam files for svviz and manual curation.

### Progression of GIAB SV Benchmark Versions

Several draft SV benchmark sets were developed and evaluated by the GIAB community, and feedback from end users and new technologies and SV callers were used to improve each subsequent version. A description of each version is below:

1. v0.2.0 included only deletions ≥20 bp, clustering was performed based only on overlap and size, and the final benchmark was a bed file with deleted regions supported by more than one technology.
2. Users of v0.2.0 requested sequence-resolved calls including insertions, so v0.3.0b included sequence-resolved deletions and insertions ≥20 bp, sequence-based clustering was performed by SVanalyzer (and experimentally by breakpoint using SURVIVOR in v0.3.0a), and the final v0.3.0b benchmark was a vcf file with sequence-resolved insertions and deletions supported by more than one technology. For v0.3.0b, only callsets with sequence-resolved calls were used. This version is under ftp://ftp-trace.ncbi.nlm.nih.gov/giab/ftp/data/AshkenazimTrio/analysis/NIST_UnionSVs_05092017/Preliminary_Integrations_v0.3.0/
3. Users of v0.3.0b found that few large insertions were included due to the requirement of support from multiple technologies, and only long reads discovered large insertions. In addition, some errors in v0.3.0b resulted from short and linked read mis-assemblies. Therefore, v0.4.0 used the same input callsets and clustering methods as v0.3.0b, and used svviz to evaluate the support for each variant by short, linked, and long reads, as well as the genotype of the SV in HG002. Calls were included in v0.4.0 if they had consistent genotypes even if they were discovered by only one technology, and calls supported by Bionano or Nabsys were also included. This version is under ftp://ftp-trace.ncbi.nlm.nih.gov/giab/ftp/data/AshkenazimTrio/analysis/NIST_UnionSVs_05092017/Preliminary_Integrations_v0.4.0/
4. Users of v0.4.0 reported that very large insertions, particularly LINEs, were still missing from the benchmark. Therefore, several improvements were made in v0.5.0:
  a. For the input callsets, especially to improve calling of large insertions, we have added:
    i. Updated sequence-resolved svanalyzer calls from assemblies, using a new version includes many more large insertions, run in discovery mode instead of targeted mode, merged calls from unscaffolded and scaffolded assemblies into single callset, and added diploid PacBio assembly from Mt Sinai.
    ii. New sequence-resolved BreakScan calls from NCBI
  b. Sites were genotyped in each member of the trio.
  c. To increase sensitivity, we excluded fewer single-tech candidate calls, requiring discovery by 4 callsets across the trio (v0.4.0 required 5 callsets in an individual)
  d. To output a heterozygous or homozygous variant GT for an individual, we required that the call was discovered in that individual (BioNano included). This eliminated some cases where the variant predicted for an individual was significantly different from the true variant in that individual.
  e. We included svviz analysis with PacBio bam separated by haplotype using whatshap with 10X variants, and we output local phasing from PacBio and 10X when available.
  f. This version is under ftp://ftp-trace.ncbi.nlm.nih.gov/giab/ftp/data/AshkenazimTrio/analysis/NIST_UnionSVs_12122017/
5. Users of v0.5.0 reported that it was useful for assessment of FNs, but not FPs, and that about a third of the v0.5.0 SVs were potentially complex or inaccurate, since they were within 1000 bp of another v0.5.0 SV. Therefore, in v0.6 we separated the calls into 2 tiers: (1) isolated, sequence-resolved SVs and (2) regions with at least one likely SV but it is complex or we were unable to determine a consensus sequence change. We also created the first SV benchmark bed file, defining regions in which the Tier 1 callset should be comprehensive, so that any extra variants detected by a method should be false positives. We also focused this benchmark to include only calls ≥50 bp and only calls in HG002, though genotypes for the parents are provided as annotations. The methods used to create v0.6 are described in this manuscript. This version is under ftp://ftp-trace.ncbi.nlm.nih.gov/giab/ftp/data/AshkenazimTrio/analysis/NIST_SVs_Integration_v0.6/

### Illumina-based SV Discovery Callsets

#### Cortex

This callset, generated jointly for the trio, used cortex^48^ (version 1.0.5.21, code at http://cortexassembler.sourceforge.net/index_cortex_var.html) with default parameters and Illumina HiSeq 300x 2×150 bp data for the AJ trio. Only variants with “PASS” status from the raw callset were included. Sites with Mendelian inconsistencies were identified and removed (47048 sites). Sites with mislabeling were also corrected (526 sites). Total variant count was 3130512, including 2780684 SNPs, 164402 deletions, 150560 insertions, 29 INV_INDEL, and 34837 COMPLEX variants, and the output VCF is under: ftp://ftp-trace.ncbi.nlm.nih.gov/giab/ftp/data/AshkenazimTrio/analysis/NCBI_IlluminaHiSeq300X_cortex_09042015/.

The input fastqs are under:

ftp://ftp-trace.ncbi.nlm.nih.gov/giab/ftp/data/AshkenazimTrio/HG002_NA24385_son/NIST_HiSeq_HG002_Homogeneity-10953946/HG002_HiSeq300x_fastq/

ftp://ftp-trace.ncbi.nlm.nih.gov/giab/ftp/data/AshkenazimTrio/HG003_NA24149_father/NIST_HiSeq_HG003_Homogeneity-12389378/HG003_HiSeq300x_fastq/

ftp://ftp-trace.ncbi.nlm.nih.gov/giab/ftp/data/AshkenazimTrio/HG004_NA24143_mother/NIST_HiSeq_HG004_Homogeneity-14572558/HG004_HiSeq300x_fastq/

#### Manta

These callsets, generated independently for each individual in the trio, used manta^49^ (version 0.27.1, code at https://github.com/Illumina/manta) with default parameters and Illumina HiSeq 30x (downsampled by read group) 2×150bp data for the AJ trio mapped by BWA MEM v1.5.0 against the hs37d5 reference genome. The output VCFs are at: ftp://ftp-trace.ncbi.nlm.nih.gov/giab/ftp/data/AshkenazimTrio/analysis/DNAnexus_AndrewC_Illumina_Callers_Sep2016/HG002/HG002.140528_D00360_0018_AH8VC6ADXX.realigned.recalibrated.manta.diploidSV.vcf

ftp://ftp-trace.ncbi.nlm.nih.gov/giab/ftp/data/AshkenazimTrio/analysis/DNAnexus_AndrewC_Illumina_Callers_Sep2016/HG003/HG003.140701_D00360_0032_AHA0KGADXX.realigned.recalibrated.manta.diploidSV.vcf

ftp://ftp-trace.ncbi.nlm.nih.gov/giab/ftp/data/AshkenazimTrio/analysis/DNAnexus_AndrewC_Illumina_Callers_Sep2016/HG004/HG004.140818_D00360_0046_AHA5R5ADXX.realigned.recalibrated.manta.diploidSV.vcf

The input fastqs, each downsampled to 30x, are under:

ftp://ftp-trace.ncbi.nlm.nih.gov/giab/ftp/data/AshkenazimTrio/HG002_NA24385_son/NIST_HiSeq_HG002_Homogeneity-10953946/HG002_HiSeq300x_fastq/

ftp://ftp-trace.ncbi.nlm.nih.gov/giab/ftp/data/AshkenazimTrio/HG003_NA24149_father/NIST_HiSeq_HG003_Homogeneity-12389378/HG003_HiSeq300x_fastq/

ftp://ftp-trace.ncbi.nlm.nih.gov/giab/ftp/data/AshkenazimTrio/HG004_NA24143_mother/NIST_HiSeq_HG004_Homogeneity-14572558/HG004_HiSeq300x_fastq/

#### GATK HaplotypeCaller

These callsets, generated independently for each individual in the trio, used GATK HaplotypeCaller^50^ (version 3.5, code at https://hub.docker.com/r/broadinstitute/gatk3) with high-sensitivity settings from Illumina HiSeq 300x 2×150 bp data for the AJ trio. Specifically, special options were ‘-stand_call_conf 2-stand_emit_conf 2 -A BaseQualityRankSumTest -A ClippingRankSumTest -A Coverage -A FisherStrand -A LowMQ -A RMSMappingQuality -A ReadPosRankSumTest -A StrandOddsRatio -A HomopolymerRun -A TandemRepeatAnnotator’. The gVCF output was converted to variant call format (VCF) using GATK Genotype gVCFs for each sample independently. The output VCFs were filtered to calls >19bp in size and are at:

ftp://ftp-trace.ncbi.nlm.nih.gov/giab/ftp/release/AshkenazimTrio/HG002_NA24385_son/NISTv3.3.2/GRCh37/supplementaryFiles/inputvcfsandbeds/HG002_GRCh37_CHROM1-MT_novoalign_Ilmn150bp300X_GATKHC.vcf.gz

ftp://ftp-trace.ncbi.nlm.nih.gov/giab/ftp/release/AshkenazimTrio/HG003_NA24149_father/NISTv3.3.2/GRCh37/supplementaryFiles/inputvcfsandbeds/HG003_GRCh37_CHROM1-Y_novoalign_Ilmn150bp300X_GATKHC.vcf.gz

ftp://ftp-trace.ncbi.nlm.nih.gov/giab/ftp/release/AshkenazimTrio/HG004_NA24143_mother/NISTv3.3.2/GRCh37/supplementaryFiles/inputvcfsandbeds/HG004_GRCh37_CHROM1-MT_novoalign_Ilmn150bp300X_GATKHC.vcf.gz

The input bam files are under:

ftp://ftp-trace.ncbi.nlm.nih.gov/giab/ftp/data/AshkenazimTrio/HG002_NA24385_son/NIST_HiSeq_HG002_Homogeneity-10953946/NHGRI_Illumina300X_AJtrio_novoalign_bams/

ftp://ftp-trace.ncbi.nlm.nih.gov/giab/ftp/data/AshkenazimTrio/HG003_NA24149_father/NIST_HiSeq_HG003_Homogeneity-12389378/NHGRI_Illumina300X_AJtrio_novoalign_bams/

ftp://ftp-trace.ncbi.nlm.nih.gov/giab/ftp/data/AshkenazimTrio/HG004_NA24143_mother/NIST_HiSeq_HG004_Homogeneity-14572558/NHGRI_Illumina300X_AJtrio_novoalign_bams/

#### Freebayes

These callsets, generated independently for each individual in the trio, used freebayes^51^ (version 0.9.20, code at https://github.com/ekg/freebayes) with high-sensitivity settings from Illumina HiSeq 300x 2×150bp data for the AJ trio. Specifically, special options were ‘-F 0.05 -m 0 --genotype-qualities’. The output VCFs were filtered to calls >19bp in size and are at:

ftp://ftp-trace.ncbi.nlm.nih.gov/giab/ftp/release/AshkenazimTrio/HG002_NA24385_son/NISTv3.3.2/GRCh37/supplementaryFiles/inputvcfsandbeds/HG002_GRCh37_CHROM1-MT_novoalign_Ilmn150bp300X_FB.vcf

ftp://ftp-trace.ncbi.nlm.nih.gov/giab/ftp/release/AshkenazimTrio/HG003_NA24149_father/NISTv3.3.2/GRCh37/supplementaryFiles/inputvcfsandbeds/HG003_GRCh37_CHROM1-Y_novoalign_Ilmn150bp300X_FB.vcf.gz

ftp://ftp-trace.ncbi.nlm.nih.gov/giab/ftp/release/AshkenazimTrio/HG004_NA24143_mother/NISTv3.3.2/GRCh37/supplementaryFiles/inputvcfsandbeds/HG004_GRCh37_CHROM1-MT_novoalign_Ilmn150bp300X_FB.vcf.gz

The input bam files are under:

ftp://ftp-trace.ncbi.nlm.nih.gov/giab/ftp/data/AshkenazimTrio/HG002_NA24385_son/NIST_HiSeq_HG002_Homogeneity-10953946/NHGRI_Illumina300X_AJtrio_novoalign_bams/

ftp://ftp-trace.ncbi.nlm.nih.gov/giab/ftp/data/AshkenazimTrio/HG003_NA24149_father/NIST_HiSeq_HG003_Homogeneity-12389378/NHGRI_Illumina300X_AJtrio_novoalign_bams/

ftp://ftp-trace.ncbi.nlm.nih.gov/giab/ftp/data/AshkenazimTrio/HG004_NA24143_mother/NIST_HiSeq_HG004_Homogeneity-14572558/NHGRI_Illumina300X_AJtrio_novoalign_bams/

#### FermiKit for 150 bp and 250 bp Illumina datasets

These callsets, generated independently for each individual in the trio, used fermikit^52^ (version 6fc8bbb3, code at https://github.com/lh3/fermikit in precisionFDA app at https://precision.fda.gov/apps/app-BvJPP100469368x7QvJkKG9Y-1) with default settings from Illumina HiSeq 50x (downsampled to two flow cells) 2×150 bp data and from Illumina HiSeq 45x 2×250 bp data for the AJ trio. Specifically, commands were ‘/fermi.kit/fermi2.pl unitig -s3g -t$(nproc) -l$(read length)-p genome reads.fq.gz > genome.mak’, ‘make -f genome.mak’, and ‘fermi.kit/run-calling -t$(nproc) bwa-indexed-ref.fa genome.mag.gz | sh’. The output VCFs were filtered to calls >19bp in size and are under: ftp://ftp-trace.ncbi.nlm.nih.gov/giab/ftp/data/AshkenazimTrio/analysis/DNAnexus_fermikit_160505/

The input fastqs are under:

ftp://ftp-trace.ncbi.nlm.nih.gov/giab/ftp/data/AshkenazimTrio/HG002_NA24385_son/NIST_HiSeq_HG002_Homogeneity-10953946/HG002_HiSeq300x_fastq/

ftp://ftp-trace.ncbi.nlm.nih.gov/giab/ftp/data/AshkenazimTrio/HG003_NA24149_father/NIST_HiSeq_HG003_Homogeneity-12389378/HG003_HiSeq300x_fastq/

ftp://ftp-trace.ncbi.nlm.nih.gov/giab/ftp/data/AshkenazimTrio/HG004_NA24143_mother/NIST_HiSeq_HG004_Homogeneity-14572558/HG004_HiSeq300x_fastq/

#### MetaSV

These callsets, generated independently for each individual in the trio, used MetaSV^53^ (version 0.5, code at https://github.com/bioinform/metasv) with default settings from Illumina HiSeq 60x 2×150 bp data for the AJ trio. Specifically, special options were ‘--boost_sc --filter_gaps --keep_standard_contigs--isize_mean 550 --isize_sd 145 --svs_to_assemble INS INV DEL DUP --svs_to_softclip INS INV DEL DUP --svs_to_report INS INV DEL DUP --max_ins_cov_frac --min_support_frac_ins 0.015 --min_support_ins 15 --max_ins_intervals 24000 --mean_read_length 150 --mean_read_coverage 60 --age_window 50’. Results from BreakSeq, BreakDancer, Pindel, and CNVnator were used as inputs into MetaSV. The output VCFs were filtered to PASS calls and are under:

ftp://ftp-trace.ncbi.nlm.nih.gov/giab/ftp/data/AshkenazimTrio/analysis/BINA_Roche_MetaSV_10142016

The input bam files are under:

ftp://ftp-trace.ncbi.nlm.nih.gov/giab/ftp/data/AshkenazimTrio/HG002_NA24385_son/NIST_HiSeq_HG002_Homogeneity-10953946/NHGRI_Illumina300X_AJtrio_novoalign_bams/

ftp://ftp-trace.ncbi.nlm.nih.gov/giab/ftp/data/AshkenazimTrio/HG003_NA24149_father/NIST_HiSeq_HG003_Homogeneity-12389378/NHGRI_Illumina300X_AJtrio_novoalign_bams/

ftp://ftp-trace.ncbi.nlm.nih.gov/giab/ftp/data/AshkenazimTrio/HG004_NA24143_mother/NIST_HiSeq_HG004_Homogeneity-14572558/NHGRI_Illumina300X_AJtrio_novoalign_bams/

#### TNscope

These callsets, generated for HG002 only, used TNscope^54^ (version 201704, from https://www.sentieon.com/) with default settings from Illumina HiSeq 300x 2×150 bp data for the AJ son. Filters removed sites with a total depth of greater than or equal to 230 (calculated as the sum of the sample AD), QUAL less than or equal to 52, or a faction of reads support the alternate allele less than

0.03. A script was used to convert the breakpoints produced by TNscope to a sequence-resolved ref/alt format for integration with the NIST callsets. The script used the orientation and size of the breakpoint to classify the breakpoint as either a deletion, duplication or inversion. The output VCFs were filtered to calls >19bp in size and are at:

ftp://ftp-trace.ncbi.nlm.nih.gov/giab/ftp/data/AshkenazimTrio/analysis/Sentieon_tnscope_05052017/HG002_300x_tnscope_hq_altallele_head.vcf.gz

The input fastqs are under:

ftp://ftp-trace.ncbi.nlm.nih.gov/giab/ftp/data/AshkenazimTrio/HG002_NA24385_son/NIST_HiSeq_HG002_Homogeneity-10953946/HG002_HiSeq300x_fastq/

#### Scalpel

These callsets, generated independently for each individual in the trio, used scalpel^55^ (version 0.4.1 beta, code at http://scalpel.sourceforge.net/) with default settings and window size 600 from Illumina HiSeq 300x 2×150 bp data for the AJ trio. The output VCFs were filtered to calls >19 bp in size and are under: ftp://ftp-trace.ncbi.nlm.nih.gov/giab/ftp/data/AshkenazimTrio/analysis/BU_HiSeq300x_scalpel_v0.4.1_04202017/

The input bam files are under:

ftp://ftp-trace.ncbi.nlm.nih.gov/giab/ftp/data/AshkenazimTrio/HG002_NA24385_son/NIST_HiSeq_HG002_Homogeneity-10953946/NHGRI_Illumina300X_AJtrio_novoalign_bams/

ftp://ftp-trace.ncbi.nlm.nih.gov/giab/ftp/data/AshkenazimTrio/HG003_NA24149_father/NIST_HiSeq_HG003_Homogeneity-12389378/NHGRI_Illumina300X_AJtrio_novoalign_bams/

ftp://ftp-trace.ncbi.nlm.nih.gov/giab/ftp/data/AshkenazimTrio/HG004_NA24143_mother/NIST_HiSeq_HG004_Homogeneity-14572558/NHGRI_Illumina300X_AJtrio_novoalign_bams/

#### SvABA

These callsets, generated jointly for the trio, used SvABA^35^ (version 0.2.1, code at https://github.com/walaj/svaba) with default settings from Illumina HiSeq 300x 2×150 bp data for the AJ trio. SvABA de-novo indel and SV calls were made with the proband BAM as the primary BAM and the parent BAMs as controls (-t <proband> -n <maternal> -n <paternal>). dbSNP v138 was used as input to the -D flag to increase confidence that de novo variants were not false negative variants from the controls. SvABA calls are produced using the breakend (BND) format, and were converted to DEL format by selecting those SVs with a +, - orientation pair, indicating a likely deletion. The variants are captured in both an SV VCF using the BND format for larger SVs, and an indel VCF for smaller SVs (< ≅100 bp).

The output VCFs were filtered to calls >19 bp in size and are under:

ftp://ftp-trace.ncbi.nlm.nih.gov/giab/ftp/data/AshkenazimTrio/analysis/Broad_svaba_05052017/ The input bam files are under:

ftp://ftp-trace.ncbi.nlm.nih.gov/giab/ftp/data/AshkenazimTrio/HG002_NA24385_son/NIST_HiSeq_HG002_Homogeneity-10953946/NHGRI_Illumina300X_AJtrio_novoalign_bams/

ftp://ftp-trace.ncbi.nlm.nih.gov/giab/ftp/data/AshkenazimTrio/HG003_NA24149_father/NIST_HiSeq_HG003_Homogeneity-12389378/NHGRI_Illumina300X_AJtrio_novoalign_bams/

ftp://ftp-trace.ncbi.nlm.nih.gov/giab/ftp/data/AshkenazimTrio/HG004_NA24143_mother/NIST_HiSeq_HG004_Homogeneity-14572558/NHGRI_Illumina300X_AJtrio_novoalign_bams/

#### Krunch

These callsets, generated independently for each individual in the trio used a method under development called Krunch (code at https://github.com/hansenlo/SeqDiff) from Illumina HiSeq 300x 2×150 bp data for the AJ trio. Krunch is a method developed to call variants that allows for direct comparison of sequence libraries with a reference genome or to each other without requiring the alignment of reads to a reference genome. The method is based on comparative indexing of DNA kmers unique to one read library compared to the other or to the reference genome. This identifies reads that share the same variant because they share the same unique kmers(s). We then assemble reads containing the same set of unique words into a contig, and align the contig or edges of the contig to the reference genome, allowing us to accurately call the variant type and position. This approach detects single nucleotide polymorphism (SNPs), small indels, medium and large structural variants, both germline and somatic. The output VCFs were filtered to calls >19 bp in size and are under:

ftp://ftp-trace.ncbi.nlm.nih.gov/giab/ftp/data/AshkenazimTrio/analysis/Stanford_Krunch_05052017/

The input fastqs are under:

ftp://ftp-trace.ncbi.nlm.nih.gov/giab/ftp/data/AshkenazimTrio/HG002_NA24385_son/NIST_HiSeq_HG002_Homogeneity-10953946/HG002_HiSeq300x_fastq/

ftp://ftp-trace.ncbi.nlm.nih.gov/giab/ftp/data/AshkenazimTrio/HG003_NA24149_father/NIST_HiSeq_HG003_Homogeneity-12389378/HG003_HiSeq300x_fastq/

ftp://ftp-trace.ncbi.nlm.nih.gov/giab/ftp/data/AshkenazimTrio/HG004_NA24143_mother/NIST_HiSeq_HG004_Homogeneity-14572558/HG004_HiSeq300x_fastq/

#### Spiral Genetics Anchored Assembly

These callsets, generated independently for each individual in the trio, used the Spiral Genetics Anchored Assembly variant caller (version May 2015, from https://www.spiralgenetics.com/) with high-sensitivity settings from Illumina HiSeq 50x (downsampled by run) 2×150bp data for the AJ trio. Specifically, the commands were ‘spiral kmerize $sample ${sample}kmers ${sample}kmer_quality_report’, ‘spiral correct_reads $sample ${sample}kmers ${sample}corrected_reads --min-kmer-score 8’, and ‘spiral find_variants ${sample}corrected_reads hg19 ${sample}variants’. The output VCFs were filtered to sequence-resolved (not breakend/BND) calls >19bp in size and are under:

ftp://ftp-trace.ncbi.nlm.nih.gov/giab/ftp/data/AshkenazimTrio/analysis/NIST_Stanford_HiSeq300x_SpiralGenetics_vcf_06042015/

The input fastqs (only run 6 for HG002 and HG004 and only run 3 for HG003) are under:

ftp://ftp-trace.ncbi.nlm.nih.gov/giab/ftp/data/AshkenazimTrio/HG002_NA24385_son/NIST_HiSeq_HG002_Homogeneity-10953946/HG002_HiSeq300x_fastq/

ftp://ftp-trace.ncbi.nlm.nih.gov/giab/ftp/data/AshkenazimTrio/HG003_NA24149_father/NIST_HiSeq_HG003_Homogeneity-12389378/HG003_HiSeq300x_fastq/

ftp://ftp-trace.ncbi.nlm.nih.gov/giab/ftp/data/AshkenazimTrio/HG004_NA24143_mother/NIST_HiSeq_HG004_Homogeneity-14572558/HG004_HiSeq300x_fastq/

#### Spiral Genetics BioGraph Refinement

This callset, generated only for HG002, used the Spiral Genetics BioGraph variant caller (version 1.1, from https://www.spiralgenetics.com/) taking in the union of all variant calls >19bp generated prior to November 11, 2016. The output VCF is under:

ftp://ftp-trace.ncbi.nlm.nih.gov/giab/ftp/data/AshkenazimTrio/analysis/Spiral_AJTrio_v1.1_01312017/ The following process was used to (1) Unite a set of GIAB Variants and unite evaluate calls using Spiral force calling. (2) Create a useful database that links all relevant information and produce metrics summarizing the results. All steps in the procedure are coded in Workflow.py. Detailed information is in Supplementary Note 1.

#### Seven Bridges Graph Refinement

This callset, generated independently for each individual in the trio, used the Seven Bridges Graph Aligner^56^ and GATK HaplotypeCaller^50^ (version 3.5, from https://hub.docker.com/r/broadinstitute/gatk3) taking in the union of all variant calls >19bp generated prior to April 14, 2017. Calls are based on alignments produced by the Seven Bridges Graph aligner, using the NIST Union SV callset 170414 from all 3 members of the trio as the contents of the reference graph. Calls are made by GATK HaplotypeCaller by explicitly forcing calls in a wide region around the putative variant sites. The output VCFs are under:

ftp://ftp-trace.ncbi.nlm.nih.gov/giab/ftp/data/AshkenazimTrio/analysis/SevenBridges_GraphGATKRefine_05052017/

### 10x Genomics-based SV Discovery Callsets

#### LongRanger

These callsets, generated independently for each individual in the trio, used LongRanger^12^ (version 2.1, code at https://github.com/10XGenomics/longranger) with default parameters and 10x Genomics data for the AJ trio (86x, 36x, and 47x coverage for HG002, HG003, and HG004, respectively). Indels >19bp from $_phased_variants.vcf.gz and large deletions from $_deletions.vcf.gz were converted into sequence-resolved vcf format. Vcf and bam files for each genome are under: ftp://ftp-trace.ncbi.nlm.nih.gov/giab/ftp/data/AshkenazimTrio/analysis/10XGenomics_ChromiumGenome_LongRanger2.1_09302016/

### Complete Genomics-based SV Discovery Callsets

#### CGATools

These callsets, generated independently for each individual in the trio, used CGATools (version 1.8.0, code at http://cgatools.sourceforge.net) with default parameters and Complete Genomics data for the AJ trio (~100x coverage for each genome). Only indels >19bp from the vcfBeta were used since SV calls were not in sequence-resolved format. vcfBeta files for each genome are at:

ftp://ftp-trace.ncbi.nlm.nih.gov/giab/ftp/data/AshkenazimTrio/analysis/CompleteGenomics_RefMaterial_SmallVariants_CGAtools_08082014/son_NA24385_GS000037263-ASM/ASM/vcfBeta-GS000037263-ASM.vcf.bz2

ftp://ftp-trace.ncbi.nlm.nih.gov/giab/ftp/data/AshkenazimTrio/analysis/CompleteGenomics_RefMaterial_SmallVariants_CGAtools_08082014/dad_NA24149_GS000037264-ASM/ASM/vcfBeta-GS000037264-ASM.vcf.bz2

ftp://ftp-trace.ncbi.nlm.nih.gov/giab/ftp/data/AshkenazimTrio/analysis/CompleteGenomics_RefMaterial_SmallVariants_CGAtools_08082014/mom_NA24143_GS000037262-ASM/ASM/vcfBeta-GS000037262-ASM.vcf.bz2

Raw data are under:

ftp://ftp-trace.ncbi.nlm.nih.gov/giab/ftp/data/AshkenazimTrio/HG002_NA24385_son/CompleteGenomics_normal_RMDNA/

ftp://ftp-trace.ncbi.nlm.nih.gov/giab/ftp/data/AshkenazimTrio/HG003_NA24149_father/CompleteGenomics_normal_RMDNA/

ftp://ftp-trace.ncbi.nlm.nih.gov/giab/ftp/data/AshkenazimTrio/HG004_NA24143_mother/CompleteGenomics_normal_RMDNA/

### PacBio-based SV Discovery Callsets

#### pbsv

These callsets, generated independently for each individual in the trio, used pbsv (version v0.1-prerelease, code at https://github.com/PacificBiosciences/pbsv) with default parameters and continuous long read PacBio data for the AJ trio aligned with NGM-LR 0.2.4 ^15^ to the hs37d5 reference (ftp://ftp.1000genomes.ebi.ac.uk/vol1/ftp/technical/reference/phase2_reference_assembly_sequence/README_human_reference_20110707). Duplicate alignments were marked with “pbsvutil markduplicates” and alignments were chained with “pbsvutil chain” with default parameters. For HG003 and HG004, at least 2 reads and 20% of reads were required. For HG002, at least 3 reads and 20% of reads were required. Only reads with MAPQ ≥ 20 were considered. VCF files for each genome are under:

ftp://ftp-trace.ncbi.nlm.nih.gov/giab/ftp/data/AshkenazimTrio/analysis/PacBio_pbsv_05052017/

Fastq files for each genome are under:

ftp://ftp-trace.ncbi.nlm.nih.gov/giab/ftp/data/AshkenazimTrio/HG002_NA24385_son/PacBio_MtSinai_NIST/

ftp://ftp-trace.ncbi.nlm.nih.gov/giab/ftp/data/AshkenazimTrio/HG003_NA24149_father/PacBio_MtSinai_NIST/

ftp://ftp-trace.ncbi.nlm.nih.gov/giab/ftp/data/AshkenazimTrio/HG004_NA24143_mother/PacBio_MtSinai_NIST/

### Multi-technology-based SV Discovery Callsets

#### HySA

These callsets, generated independently for each individual in the trio, used HySA^57^ (commit ID eee31f6, code at https://bitbucket.org/xianfan/hybridassemblysv/overview) with default parameters and merged Illumina HiSeq 300x 2×150bp data and continuous long read PacBio data for the AJ trio. Post filtering includes only one step: removing all calls with just one Illumina read as the support. Vcf files for each genome are under: ftp://ftp-trace.ncbi.nlm.nih.gov/giab/ftp/data/AshkenazimTrio/analysis/MDAnderson_HySA_05052017/

Fastq files for each genome are under:

ftp://ftp-trace.ncbi.nlm.nih.gov/giab/ftp/data/AshkenazimTrio/HG002_NA24385_son/NIST_HiSeq_HG002_Homogeneity-10953946/HG002_HiSeq300x_fastq/

ftp://ftp-trace.ncbi.nlm.nih.gov/giab/ftp/data/AshkenazimTrio/HG003_NA24149_father/NIST_HiSeq_HG003_Homogeneity-12389378/HG003_HiSeq300x_fastq/

ftp://ftp-trace.ncbi.nlm.nih.gov/giab/ftp/data/AshkenazimTrio/HG004_NA24143_mother/NIST_HiSeq_HG004_Homogeneity-14572558/HG004_HiSeq300x_fastq/

ftp://ftp-trace.ncbi.nlm.nih.gov/giab/ftp/data/AshkenazimTrio/HG002_NA24385_son/PacBio_MtSinai_NIST/

ftp://ftp-trace.ncbi.nlm.nih.gov/giab/ftp/data/AshkenazimTrio/HG003_NA24149_father/PacBio_MtSinai_NIST/

ftp://ftp-trace.ncbi.nlm.nih.gov/giab/ftp/data/AshkenazimTrio/HG004_NA24143_mother/PacBio_MtSinai_NIST/

#### BreakScan

BreakScan is a kmer-based structural variation discovery method, which models insertion and deletion events in reference sequence with breakpoint junctions observed in NGS reads, then subsequently compiles evidence for those junctions from the sequencing reads generated by multiple platforms such as Illumina, 10X Genomics, and Pacific Biosciences. The candidate structural variants are generated from event models and ranked by their supporting evidence. These callsets, including 2,918 deletions and 2,193 insertions, generated only for HG002, used BreakScan (https://github.com/chunlinxiao/BreakScan) with default parameters and reads from Illumina HiSeq 300x 2×150bp data, 10x Genomics data, and error-corrected continuous long read PacBio data for HG002. Only variants supported by at least two technologies are included. VCF files for insertions and deletions are under:

ftp://ftp-trace.ncbi.nlm.nih.gov/giab/ftp/data/AshkenazimTrio/analysis/NCBI_BreakScan_12072017_v1.1/

Input fastq files for each technology are under:

ftp://ftp-trace.ncbi.nlm.nih.gov/giab/ftp/data/AshkenazimTrio/HG002_NA24385_son/NIST_HiSeq_HG002_Homogeneity-10953946/HG002_HiSeq300x_fastq/,

ftp://ftp-trace.ncbi.nlm.nih.gov/giab/ftp/data/AshkenazimTrio/analysis/10XGenomics_ChromiumGenome_LongRanger2.1_09302016/

ftp://ftp-trace.ncbi.nlm.nih.gov/giab/ftp/data/AshkenazimTrio/HG002_NA24385_son/PacBio_MtSinai_NIST/

### Global de novo assembly-based SV Discovery Callsets

#### SVrefine

These callsets, generated independently for each individual in the trio, used SVRefine (version 0.2, code at https://github.com/nhansen/SVanalyzer) with default parameters and from a variety of global de novo assemblies from different technologies and assembly methods. For assemblies with both unscaffolded fasta files and fasta files scaffolded with Dovetail, we merged calls from unscaffolded assemblies with their Dovetail-scaffolded counterparts using SVmerge.pl and SVcluster.pl from the SVanalyzer, with the commands ‘SVmerge.pl --ref $REF --vcf $UNIONVCF > $DISTFILE’, ‘gunzip -c $VCF | grep -v ‘#’ | awk ‘{print $3}’ > $IDFILE’, and ‘SVcluster.pl --ids $IDFILE --dist $DISTFILE --relshift 0 --relsizediff 0 --reldist 0 --vcf $VCF > $CLUSTERFILE’. Specifically, HG2_MHAP_plus_Dovetail.clustered.0.0.0.vcf.gz is a merge of HG2_Dovetail_MHAP.l100c500.vcf.gz and HG2_PBcR_MHAP.l100c500.vcf.gz, HG2_Falcon_plus_Dovetail.clustered.0.0.0.vcf.gz is a merge of HG2_Dovetail_Falcon.l100c500.vcf.gz and HG2_p_and_a_ctg.l100c500.vcf.gz, and nd HG2_DISCOVAR_plus_Dovetail.clustered.0.0.0.vcf.gz is a merge of HG2_Dovetail_DISCOVAR.l100c500.vcf.gz and HG2_DISCOVAR250bp.l100c500.vcf.gz.

De novo assemblies used as inputs to SVRefine were:

PacBio-only:

ftp://ftp-trace.ncbi.nlm.nih.gov/giab/ftp/data/AshkenazimTrio/analysis/UMD_PacBio_Assembly_CA8.3_08252015/

ftp://ftp-trace.ncbi.nlm.nih.gov/giab/ftp/data/AshkenazimTrio/analysis/MtSinai_PacBio_Assembly_falcon_03282016/

Illumina-only:

ftp://ftp-trace.ncbi.nlm.nih.gov/giab/ftp/data/AshkenazimTrio/analysis/TAMU_NIST_Illumina_2x250bps_DISCOVAR_Assemblies_09162016/

Dovetail-scaffolded assemblies from PacBio and Illumina:

ftp://ftp-trace.ncbi.nlm.nih.gov/giab/ftp/data/AshkenazimTrio/analysis/Dovetail_HiRiseScaffolding_10142016/10x

Genomics:

ftp://ftp-trace.ncbi.nlm.nih.gov/giab/ftp/data/AshkenazimTrio/analysis/10XGenomics_ChromiumGenome_LongRanger2.2_Supernova2.0.1_04122018/assemblies/ HG004 Illumina 2×250 paired end and 6kb mate-pair sequencing, plus 10x Genomics Chromium linked reads and Bionano optical mapping for scaffolding, using ABySS 1.9, ABySS 2.0, BCALM2, DISCOVARdenovo, Megahit, Minia, SGA, and SOAPdenovo (https://genome.cshlp.org/content/early/2017/02/23/gr.214346.116):

ftp://ftp-trace.ncbi.nlm.nih.gov/giab/ftp/data/AshkenazimTrio/analysis/BCGSC_HG004_ABySS2.0_assemblies_12082016/

Vcf files for each genome from each assembly are under:

ftp://ftp-trace.ncbi.nlm.nih.gov/giab/ftp/data/AshkenazimTrio/analysis/NHGRI_SVrefine_12062017/

#### Assemblytics

These callsets, generated independently for each individual in the trio, used assemblytics^58^ (version 1.0, code at https://github.com/MariaNattestad/Assemblytics/releases/tag/v1.0) with default parameters and from two haploid de novo assemblies from PacBio. For each genome assembly, the assembly fasta file was aligned to the reference using MUMmer (v3.23) nucmer method with the following parameters:

-maxmatch -l 100 -c 500. Assemblytics was run with default parameters (10,000 bp unique sequence anchor length) on the delta file output from nucmer. Results were transformed into VCF format using SURVIVOR^40^ and a custom script, and filtered to variants ≥ 20 bp long.

Haploid de novo assemblies used as inputs to assemblytics were fromPacBio-only:

ftp://ftp-trace.ncbi.nlm.nih.gov/giab/ftp/data/AshkenazimTrio/analysis/UMD_PacBio_Assembly_CA8.3_08252015/

ftp://ftp-trace.ncbi.nlm.nih.gov/giab/ftp/data/AshkenazimTrio/analysis/MtSinai_PacBio_Assembly_falcon_03282016/

#### MsPAC

These callsets, generated independently for each individual in the trio, used MsPAC v.e30c77e ((https://github.com/oscarlr/MsPAC) with default parameters. PacBio reads aligned to GRCh37 were downloaded from ftp://ftp-trace.ncbi.nlm.nih.gov/giab/ftp/data/AshkenazimTrio/HG002_NA24385_son/PacBio_MtSinai_NIST/MtSinai_blasr_bam_GRCh37 and phased SNPs were downloaded from

ftp://ftp-trace.ncbi.nlm.nih.gov/giab/ftp/data/AshkenazimTrio/analysis/10XGenomics_ChromiumGenome_LongRanger2.0_06202016/HG002_NA24385_son/NA24385_GRCh37.vcf.gz. Using the PacBio aligned bam files and 10X phased SNVs as input, MsPAC generated diploid assemblies and phased SVs calls.

Assembly fasta/fastq files as wells as VCF files for called SVs can found here:

ftp://ftp-trace.ncbi.nlm.nih.gov/giab/ftp/data/AshkenazimTrio/analysis/MSSM_MsPAC_SVs_assemblies_06042019/

#### Phased-SV

Haplotype-specific assemblies and associated callsets for HG002 were generated using Phased-SV (github.com/mchaisso/phasedsv) with parameters {“recall_bin”: 100, “cov_cutoff”: 3, “tr_cluster_size”: 6, “depth” : 50}. Reads were aligned to GRCh38 using blasr (github.com/mchaisso/blasr) retaining quality value information. SNP phasing from ftp://ftp-trace.ncbi.nlm.nih.gov/giab/ftp/data/AshkenazimTrio/analysis/10XGenomics_ChromiumGenome_LongRanger2.0_06202016/HG002_NA24385_son/NA24385_GRCh37.vcf.gz was used to partition reads by haplotype, and local assemblies were performed using canu.^59^ Insertion and deletion SV calls are available at ftp://ftp-trace.ncbi.nlm.nih.gov/giab/ftp/data/AshkenazimTrio/analysis/Chaisson_PacBio_smrt-sv.dip_Jun2016/

### De novo assemblies

#### PacBio Canu (haploid)

A non-diploid assembly was generated using the CA 8.3 assembly method^59^ from PacBio Continuous Long Read data from each member of the Ashkenazi Trio. The assemblies used MHAP “sensitive” parameters and PBDAGCON for consensus. All assemblies have been polished using Quiver. The assemblies are available at:

ftp://ftp-trace.ncbi.nlm.nih.gov/giab/ftp/data/AshkenazimTrio/analysis/UMD_PacBio_Assembly_CA8.3_08252015/ PacBio data used for each member of the trio is in NCBI SRA SRX1033793–SRX1033798

#### PacBio Falcon (haploid)

A non-diploid assembly was generated using the Falcon assembly method^18^ from PacBio Continuous Long Read data from each member of the Ashkenazi Trio. The assemblies are available at:

ftp://ftp-trace.ncbi.nlm.nih.gov/giab/ftp/data/AshkenazimTrio/analysis/MtSinai_PacBio_Assembly_falcon_03282016/

PacBio data used for each member of the trio is in NCBI SRA SRX1033793–SRX1033798

#### Illumina DISCOVAR (haploid)

A non-diploid assembly was generated using the DISCOVAR De Novo tool^60^ (https://software.broadinstitute.org/software/discovar/blog/) from 2×250bp Illumina sequencing data from each member of the Ashkenazi Trio. The assemblies are available at:

ftp://ftp-trace.ncbi.nlm.nih.gov/giab/ftp/data/AshkenazimTrio/analysis/TAMU_NIST_Illumina_2x250bps_DISCOVAR_Assemblies_09162016/

Illumina 2×250bp overlapping read data used for DISCOVAR is in NCBI SRA:

SRX1726837-SRX1726840, SRX1726853-SRX1726856, SRX1726860, SRX1726868, and SRX1726870 for HG002

SRX1726871-SRX1726875 and SRX1726881-SRX1726893 for HG003

SRX1726894-SRX1726928 for HG004

#### Dovetail Chicago Scaffolding of PacBio and Illumina Assemblies

Dovetail Genomics generated Chicago libraries on HG002, HG003, and HG004 and used HiRise to scaffold 3 existing assemblies for each genome:

1. PacBio Falcon: ftp://ftp-trace.ncbi.nlm.nih.gov/giab/ftp/data/AshkenazimTrio/analysis/MtSinai_PacBio_Assembly_falcon_03282016/
2. PacBio PBcR/MHAP: ftp://ftp-trace.ncbi.nlm.nih.gov/giab/ftp/data/AshkenazimTrio/analysis/UMD_PacBio_Assembly_CA8.3_08252015/
3. Illumina 2×250bp DISCOVAR: ftp://ftp-trace.ncbi.nlm.nih.gov/giab/ftp/data/AshkenazimTrio/analysis/TAMU_NIST_Illumina_2x250bps_DISCOVAR_Assemblies_09162016/

Raw reads are under

ftp://ftp-trace.ncbi.nlm.nih.gov/giab/ftp/data/AshkenazimTrio/HG.../Dovetail_ChicagoLibraties/ for each genome.

Scaffolded assembly results are under each genome

ftp://ftp-trace.ncbi.nlm.nih.gov/giab/ftp/data/AshkenazimTrio/analysis/Dovetail_HiRiseScaffolding_1014 2016, with a description of the files in manifest.txt under the hu… directory under each starting assembly. bam files with reads mapped to the assembly are under the bans directory for each genome.

### Optical and Electronic Mapping

#### Bionano

These callsets, generated independently for each individual in the trio, used Bionano Solve v3.1 (bnxinstall.com/solve/Solve3.1_08232017) with default parameters and from Bionano data generated from two enzymes BspQI and BssSI. SV calls and maps are available under:

ftp://ftp-trace.ncbi.nlm.nih.gov/giab/ftp/data/AshkenazimTrio/analysis/BioNano_haplotype_SV_BspQI_BssSI_overlapv0.3.0b_10312017/

#### Nabsys

This callset, generated for HG002, used Nabsys HD-Mapping with NPS Analysis v1.2.1922 and SV-Verify 12.0.^20^ Single molecule reads, ≥50kb were mapped to both GRCh37 and constructs representing putative deletions. Mapping results were evaluated by a set of support vector machines that had been trained on similar sized deletions in NA12878. SV-Verify tests the hypothesis that the specified number of bases, as defined by a putative deletion, are deleted in the region between the lower and upper bound probe locations, specified in the .bed file. Additional SVs (deletions, insertions) occurring within the same region will invalidate the hypothesis. SV-Verify results are under:

ftp://ftp-trace.ncbi.nlm.nih.gov/giab/ftp/data/AshkenazimTrio/analysis/Nabsys_v0.3.0bCorroboration_Sept2017/

### Callsets benchmarked against v0.6 Tier 1 benchmark set

GIAB asked for volunteers to compare their SV callsets to the v0.6 Tier 1 benchmark set with truvari, and manually curate 10 randomly selected FPs and FNs each from insertions and deletions, subset to SVs overlapping and not overlapping tandem repeats longer than 100bp (80 total variants). Potential errors identified in GIAB were further examined by NIST and the final determination about whether v0.6 was correct was made in consultation between multiple curators.

#### Illumina mapping-based Delly and Manta

Structural variants were called from Illumina HiSeq 300x 2×150bp data (previously aligned to hs37d5) using Manta (version 1.2.2, code at https://github.com/Illumina/manta) with default options and Delly (version 0.7.8, code at https://github.com/dellytools/delly) with minimum mapping quality set to 20. For Manta, all calls from diploidSV.vcf output file were filtered for the “PASS” filter field. For Delly, SVs were discovered, merged with Delly’s default values (breakpoint offset: 1000 & reciprocal overlap: 0.8) and genotyped. Output calls were filtered according to the “PASS” filter and a minimal count of alternate-supporting reads of 5. SVs in centromeres and telomeres were excluded, with the list provided with Delly developers. Calls were compared to the goldset using truvari, and manually verified in IGV.

#### Illumina mapping-based MetaSV

The same MetaSV callset described above was benchmarked against v0.6.

#### Illumina assembly-based SpiralBGA

The same Spiral BioGraph methods described above was benchmarked against v0.6

#### PacBio mapping-based pbsv

PacBio 10 kb CCS reads (ftp://ftp-trace.ncbi.nlm.nih.gov/giab/ftp/data/AshkenazimTrio/HG002_NA24385_son/PacBio_CCS_10kb/) were aligned to hs37d5 with minimap2 version 2.11-r797 (https://github.com/lh3/minimap2). ^61^ Structural variants were called with pbsv version 2.0.0 (https://github.com/PacificBiosciences/pbsv) with default parameters. Variants were evaluated against the GIAB v0.6 benchmark set using Truvari commit bb51e7575 with “--passonly --pctsim 0 -r 2000 --giabreport”. Ten randomly selected variants were evaluated in IGV for each combination of variant type (insertion, deletion); Truvari error type (false positive, false negative); and overlap with tandem repeats (yes, no). The pbsv VCF files are at: ftp://ftp-trace.ncbi.nlm.nih.gov/giab/ftp/data/AshkenazimTrio/analysis/PacBio_pbsv_05212019/

#### PacBio assembly-based SVRefine from MsPAC Diploid Assembly

The following vcfs were combined with svanalyzer, merging calls within 20% edit distance, and compared to the v0.6 Tier 1 benchmark with truvari:

ftp://ftp-trace.ncbi.nlm.nih.gov/giab/ftp/data/AshkenazimTrio/analysis/NHGRI_SVrefine_05252018/hs37d5/HG2_ORod_raw.01_0518.l100c500.no_ns.vcf.gz

ftp://ftp-trace.ncbi.nlm.nih.gov/giab/ftp/data/AshkenazimTrio/analysis/NHGRI_SVrefine_05252018/hs37d5/HG2_ORod_raw.02_0518.l100c500.no_ns.vcf.gz

These vcfs were generated from each assembled haplotype of MsPAC in the folder below using SVRefine, as described above:

ftp://ftp-trace.ncbi.nlm.nih.gov/giab/ftp/data/AshkenazimTrio/analysis/MSSM_MsPAC_SVs_assemblies_06042019/

#### 10x Genomics mapping-based LongRanger

This callset used LongRanger (version 2.2, code at https://github.com/10XGenomics/longranger) with default parameters and 10x Genomics data for HG002 (86x). Indels >19bp from

$_phased_variants.vcf.gz and large deletions from $_deletions.vcf.gz were converted into sequence-resolved vcf format. Vcf and bam files for each genome are under:

ftp://ftp-trace.ncbi.nlm.nih.gov/giab/ftp/data/AshkenazimTrio/analysis/10XGenomics_ChromiumGenome_LongRanger2.2_Supernova2.0.1_04122018/

#### Bionano Genomics

GM24385 data generated using the DLS chemistry and Bionano Saphyr system are assembled and have SVs called against hg19 using Bionano Solve v3.2.2 (bnxinstall.com/solve/Solve3.2.2_08222018)) with default parameters. The data and SV calls are under:

ftp://ftp-trace.ncbi.nlm.nih.gov/giab/ftp/data/AshkenazimTrio/analysis/BioNano_haplotype_SV_06072019

Bionano indels (>1 kbp) are overlapped with the 1557 v0.6 benchmark calls (>1 kbp) that showed “PASS” in the FILTER column; Bionano calls with 80% size concordance and 50% reciprocal position overlap are clustered to avoid duplicated counts of homozygous calls. Between the Bionano calls and the v0.6 benchmark calls, a size concordance of 50% and breakpoint precision of 5kbp are required for them to be overlapped. About 90% of the v0.6 benchmark calls overlapped with Bionano, and an additional 1 % match when summing of neighboring indels within the same Bionano markers. Only a few Bionano unique calls fall within the Tier1 region, but over a thousand Bionano unique calls fall outside of the Tier1 regions, where Bionano may be able to detect SVs in repetitive regions spanned by its ultra-long (323 kbp N50) molecules. Future work will be needed to develop robust benchmarks for complex events and very large, repetitive regions.

### Other SV Callsets

We generated additional SV callsets, which were not used in forming or evaluating the v0.6 SV benchmark set, but some were used in previous benchmark versions or are a resource for community evaluations.

*Illumina mapping-based TARDIS*

These call sets are generated jointly for the trio used TARDIS^62^ (version 1.0.4, code at https://github.com/BilkentCompGen/tardis) with default settings from Illumina HiSeq 300x and 100x 2×150bp data for the AJ and Chinese trios. TARDIS SV calls were made from the BAM files and all SVs with read pair support < 50 were filtered out. For the genomes with 100x depth of coverage, the minimum read pair support was 18. TARDIS call sets includes deletions, inversions, tandem and interspersed duplications, mobile element insertions, nuclear mitochondrial DNA insertions, and small novel insertions. Only those SVs that are supported by multiple soft-clipped reads are sequence-resolved. The output VCF and BED files are under: ftp://ftp-trace.ncbi.nlm.nih.gov/giab/ftp/data/AshkenazimTrio/analysis/BilkentUni_IlluminaHiSeq_TARDIS_mrCaNaVar_05212019/

ftp://ftp-trace.ncbi.nlm.nih.gov/giab/ftp/data/ChineseTrio/analysis/BilkentUni_IlluminaHiSeq_TARDIS_mrCaNaVar_05212019/

#### Illumina mapping-based MrCaNaVaR

We used mrCaNaVaR tool^63^ with default parameters to characterize large (>10 Kb) segmental duplications and deletions, and calculate genic copy numbers. The mrCaNaVaR tool is a reimplementation of an earlier read depth based algorithm designed to detect segmental duplications. Briefly, we remapped the Illumina reads to the repeat-masked reference genome assembly, and identified regions of read depth higher than the genome average (specifically 3 standard deviations above average) after GC% error correction. The output VCF and BED files are under:

ftp://ftp-trace.ncbi.nlm.nih.gov/giab/ftp/data/AshkenazimTrio/analysis/BilkentUni_IlluminaHiSeq_TARDIS_mrCaNaVar_05212019/

ftp://ftp-trace.ncbi.nlm.nih.gov/giab/ftp/data/ChineseTrio/analysis/BilkentUni_IlluminaHiSeq_TARDIS_mrCaNaVar_05212019/

#### PacBio mapping-based PALMER

We used PALMER (https://github.com/mills-lab/PALMER) to identify non-reference human-specific Long Interspersed Element-1 (L1Hs) insertions and characterize significant hallmarks of these retrotransposon insertions. PALMER firstly pre-masks aligned long-read sequences containing endogenous reference L1Hs and then searches against L1.3 (GenBank Accession: L19088) sequences to detect non-reference L1Hs insertions within the remaining unmasked sequences. After we obtained the preliminary non-reference L1Hs insertions from PALMER, error-correction and local-alignment processes were manipulated by using CANU^59^ (https://github.com/marbl/canu) and blasr (github.com/mchaisso/blasr). We run PALMER onto these error-corrected reads and obtained a high confident set of each sample. The output VCF files, including L1Hs insertion sequences and 5’ and 3’ target site duplicate sequences from PacBio data, are under:

ftp://ftp-trace.ncbi.nlm.nih.gov/giab/ftp/data/AshkenazimTrio/analysis/PacBio_PALMER_11242017/

## Supplementary Information

**Supplementary Figure 1:**
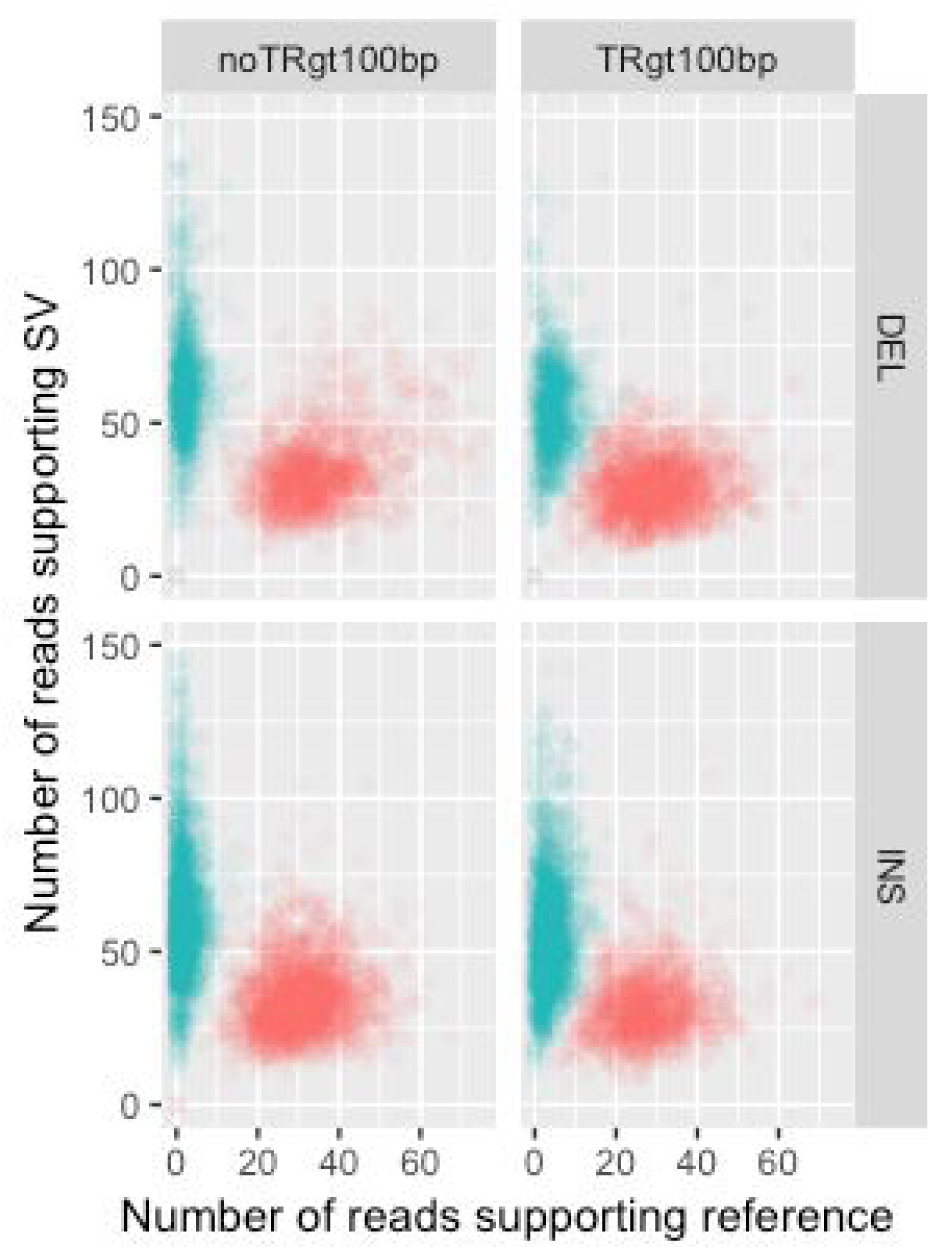
Number of long reads supporting the SV allele vs. the reference allele in the benchmark set. Variants are colored by heterozygous (red) and homozygous (blue) genotype, and are stratified into deletions (DEL) and insertions (INS), and into SVs overlapping (TRgt100bp) and not overlapping (noTRgt100bp) tandem repeats longer than 100 bp in the reference.

**Supplementary Table 1:**
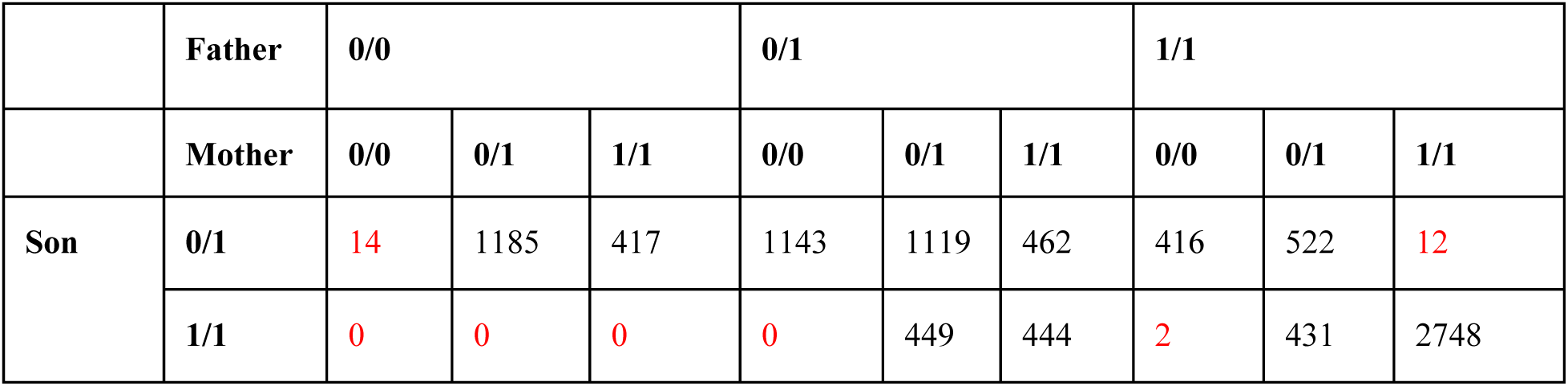
Mendelian contingency table for sites with consensus genotypes from svviz in the son, father, and mother. SVs in boxes highlighted in red violate the expected Mendelian inheritance pattern. Variants on chromosomes X and Y are excluded.

|  | Father | 0/0 |  |  | 0/1 |  |  | 1/1 |  |  |
| --- | --- | --- | --- | --- | --- | --- | --- | --- | --- | --- |
|  | Mother | 0/0 | 0/1 | 1/1 | 0/0 | 0/1 | 1/1 | 0/0 | 0/1 | 1/1 |
| Son | 0/1 | 14 | 1185 | 417 | 1143 | 1119 | 462 | 416 | 522 | 12 |
|  | 1/1 | 0 | 0 | 0 | 0 | 449 | 444 | 2 | 431 | 2748 |

***Supplementary Note 1: Spiral BioGraph Refinement Process***

Individual stages can be executed one at a time or the entire process can be run with ‘python Workflow.py all’

**## Step 1 - makeDB - Create Database**

This removes any existing database and loads the schema in DBSchema.sql into an sqlite3 database named AJTrio.db

Tables:

- GIABVariant
  - Holds a single GIAB Variant as loaded from the union of all variant calls >19bp generated prior to November 11, 2016
- Locus
  - Represents the coordinates of a set of merged GIABVariants
- LocusCalls
  - Holds which GIABVariants are within each Locus
- SpiralVariant
  - Spiral Variant that was created through force calling
- ForceMatch
  - Holds the relation between how each GIABVariant within a Locus was force called with a SpiralVariant
- Trio
  - Matches SpiralVariants across individuals that are identical - representing inheritance.
- Collection
  - Represents the relationship between the elements above. Per locus, we want to show how close any particular GIABVariant is to any Trio Call.

**## Step 2 - loadGIAB - Load GIAB Variants**

This parses the input GIABVariants and loads them into our GIAB Table.

In order to homogeneously represent all the variants in the database. Some massaging of the data had to be performed.

1. Insertions with identical starts and ends had +1 added to the end to prevent bedtools having problems sorting entries.
2. Chromosome MTT was changed to MT
3. The variant’s name entry was split by ‘_’ with properties individual, platform, program, other. However, name property was still preserved
4. Qualities of ‘.’ were translated to -1
5. The end was set with the following logic:
  1. If END was in the info, set to that
  2. else if svtype is INS - set end to start
  3. else if svtype is DEL - set end to start + svlength
  4. if the svtype is one of “BND”, “INV_INDEL”, we do not set the end coordinate
6. If end was before the start, switch the two values
7. The svtype was set with the following logic:
  1. If SVTYPE, TYPE or SV_TYPE is in info, use that value as the svtype.
  2. Else if the length of the refSeq is greater than the length of the altSeq, it’s a deletion
  3. Else if the length of the altSeq is less than the length of the refSeq, it’s an insertion
  4. If the svtype is complex - use logic of #2 and #3 above
8. The svlen was set with the following logic:
  1. If SVLEN in info, use that value
  2. else infer the size using altSeq/refSeq or the distance between start and end

**## Step 3 - filtGIAB - Filter GIAB Variants**

set ignore flags in the GIABVariants table’s removed column.

1. if variant’s start < 0 : set bad_coord
2. if svtype != ins or del : set bad_type
3. if end-start > MAXSPAN (20k) : set large_span
4. if svlen > MAXLEN (20k) : set large_size
5. If variant intersects with a reference gap : set gap type

**## Step 4 - makeLoci - Merge GIAB Variants and Create Loci** Combine the GIABVariants to create loci where we’ll be force calling. Calls were separated by svtype (only DEL and INS were considered) Used bedtools merge with book-ends distance of MERGEDIST (100)

**## Step 5 - cluster - Create Spiral force calls**

For every locus, we took a ±300bp buffer from the start and end If this was ≥ 10kb - we ignored the call.

Resulting force calls were filtered with the following conditions:

1. The variant’s annotation is not MNP, SNP, or REF
2. At least 5 reads supported the alternate
3. The variant’s length was at least 20bp
4. The variant’s annotation starts with the locus’ svtype
  1. Note: Many spiral assemblies’ remapping on a reference will be a ‘net-gain’ insertion of ‘net-loss’ deletion. For example, if 500bp is removed from the sample relative to the reference and replaced with 400bp, this is considered a 100bp DELrp - Deletion with replacement. This example variant can only match with loci with svtype DEL

Each of these force calls are recorded in the SpiralVariant table. The ForceMatch holds what SpiralVariants match to which GIABVariants with metrics.

The logic for matching a the resulting force calls from above is as follows:

1. If force calling failed (couldn’t assemble anything in the region), We add an entry for the locus that is annotated as ‘ERROR’
2. If there is only a single reference force call, We add that entry into SpiralVariant and annotate the ForceMatch as “REF”
  1. These places may generally be considered “Likely False Positives”
3. If there are no force calls passing the above filtering and more than a single reference force call, No SpiralVariant is recorded and the ForceMatch is annotated as “Unknown”
  1. This could mean a wide variety of things, but the one thing we know for sure is that none of the calls helped us get to a structural variant
4. If there is at least one force call passing the above filtering, for every GIABVariant inside the locus, we choose the one force call with the closest euclidean distance to compare to the variant. We then check if either of the following two conditions are true.
  1. There’s less than 100 euclidean distance from the force call’s start/end and the variant’s start/end
  2. There’s at least 80% reciprocal overlap between the force call and variant positions.
5. If 4.1 or 4.2 are true, we annotate the ForceCall as “Match”, if not, we annotate as “Near”. Note that in loci with multiple force calls, we only report SpiralVariants in the ForceCall table if they are the nearest to at least one variant.

